# Bio-inspired mineralization collagen induce fibrocartilage regeneration after tendon-bone injury by activating Gli1+Dkk3+ progenitor cells

**DOI:** 10.1101/2023.09.24.557863

**Authors:** Tao Zhang, Tingyun Lei, Jie Han, Ru Zhang, Weiliang Shen, Yi Liu, Youguo Liao, Yanyan Zhao, Xianzhu Zhang, Ruojin Yan, Qiulin He, Yangwu Chen, Haihua Pan, Ouyang Hongwei, Lingting Wang, Wei Yin, Zi Yin, Chen Xiao

## Abstract

A fibrocartilaginous connection between the tendon and bone, plays a critical role in transferring force from muscle to bone to enable joint movement. However, due to the high mechanical stress it experiences, the enthesis is vulnerable to injury and incapable of regenerating. The spatial relationship and functional basis of the principal components of the fibrocartilage - mineral and collagen - have not been clearly elucidated, which is a significant remaining gap in reconstructing complex architectures for promoting interface tissue regeneration. Here, using three-dimensional electron tomography imaging and high-resolution two-dimensional electron microscopy, we discover that mineral particles form a continuous cross-fibrillar phase within the fibrocartilage region. By developing a “floating mineralization” system, we fabricate a three-layer hydrogel that mimics the hierarchical nano- to micro-scale structure of tendon-bone interface (TBI). The middle layer is noteworthy for its resemblance to the nanostructure of fibrocartilage and its superior ability to induce mineralized fibrochondrogenesis *in vitro*. Based on motor function analysis, imaging diagnosis, histological staining, immunofluorescence staining, and biomechanics performance, we demonstrate that in situ transplantation of the gradient hydrogel achieved tendon-fibrocartilage-bone synchronous regeneration and result in 68% maximum mechanical recovery at 8-week postoperation. Single-cell RNA sequencing analysis reveals that a unique atlas of in situ stem/progenitor cells is generated during the TBI healing *in vivo*. Notably, the bio-inspired hydrogel microenvironment drived endogenous Gli1^+^Dkk3^+^ progenitor cells, playing a key role in TBI regeneration. Therefore, we have successfully decoded and reconstructed the nanostructure of fibrocartilage, which has great potential in TBI regeneration.

## Introduction

The tendon-bone interface (TBI), referred to as the enthesis, delineates the point of insertion where tendons and ligaments merge with the bone surface^[1]^. A fibrocartilaginous connection between the tendon and bone, plays a critical role in transferring force from muscle to bone to enable joint movement. Within the realm of sports medicine, TBI injuries, encompassing afflictions such as rotator cuff and anterior cruciate ligament tears or detachment, stand as prevalent conditions^[1]^. While the established clinical norm involves surgically reattaching tendons or ligaments to bone, this practice culminates in the formation of scar tissue devoid of fibrocartilage transition. The consequent abrupt demarcation between mechanically incongruous tendon and bone induces strain concentrations that markedly amplify the susceptibility to re-failure (>90% in some older demographics)^[2]^. Emerging as a beacon of hope for the effective restoration of TBI injuries, in situ tissue engineering demands meticulous mastery over the biophysical and biochemical stimuli inherent in biomaterials, a mastery that guides endogenous cells precisely to the injury locus. These stimuli are indispensable, serving to incite regeneration through modulation of the extracellular microenvironment or instigation of cellular reprogramming^[3]^.

The composition and structure of natural tissues offer insights into potential biomaterial design. The inorganic composition of the TBI is divided into three gradients: non-mineralized area (tendon) with 0 wt% inorganic content; moderately mineralized area (fibrocartilage) with 32 wt% inorganic content; and highly mineralized area (bone) with 65 wt% inorganic content^[4, 5]
^. Mechanically, in the direction of muscle action, the elastic modulus of tendon tissue transitions from ∼0.45 GPa to ∼20 GPa in bone tissue^[2]^. Upon observation at the nanometer scale, it was discovered that tendons are composed of non-mineralized Type I collagen fibrils. The collagen molecules (diameter = 1.5 nm) self-assemble into fibrils with a 27 nm overlap and a 40 nm gap; this configuration yields a characteristic d-spacing of 67 nm^[6]^. Among them, the channels I and II (1.5 nm high × 5.8 nm wide) and the channel III (1.5 nm high × 5.6 nm wide) formed in the gap region provide the spatial foundation for the infiltration of inorganic substances^[7]^. Bones are predominantly composed of mineralized collagen I fibrils. The precursor phase for bone formation is amorphous calcium phosphate (ACP), rather than hydroxyapatite (HAP). The ACP, stabilized by non-collagenous proteins (NCPs), is consisted of densely packed clusters of about 1 nm in size^[8, 9]^. The accumulated minerals exceed the lateral dimensions of the collagen fibrils and span across adjacent fibrils, forming continuous cross-fibrillar mineralization^[10, 11]^. The fibrocartilage is characterized by partial inorganic deposition^[12]^. This attribute bestows the ability for mobility of inorganic constituents, facilitating the dissipation of mechanical forces during conduction and endowing it with toughness under varied loadings^[13]^. Nonetheless, the interplay between fibrocartilage’s matrix principal components— mineral and collagen fibrils—remains incompletely elucidated. Notably, three-dimensional high-resolution (3D-HR) techniques, particularly focused ion beam-scanning electron microscopy (FIB-SEM), have been employed to investigate mineralized tissues, yielding comprehensive insights into their structural attributes^[11, 14]^. In this study, utilizing FIB-SEM, we observed that within the mineralized region of fibrocartilage, mineral particles do not exclusively align intrafibrillarly or extrafibrillarly; instead, they form a continuous cross-fibrillar phase. Therefore, the TBI initiate assembly from several nanoscale collagen molecules and calcium phosphate (CaP) nanoclusters, culminating in the formation of tissues characterized by gradient transitions in structure, composition, and mechanical properties.

Recently, we conducted a comprehensive review of the latest advancements in biomaterial design to TBI defects^[15]^. The utilization of organic materials such as poly(L-lactic) acid, poly(ε-caprolactone), and others presents a challenge in replicating the nanoscale features found in natural fibrils, particularly the tubular and channel structures. On the other hand, inorganic materials, such as HAP and CaP silicate, are often utilized in sizes ranging from tens to hundreds of nanometers. These materials prevent inorganic crystals penetration into the subchannels of the fibrils, thereby hindering the precise attainment of a TBI scaffold^[16–20]^. It is noteworthy that the topological structure of materials plays a crucial role in cellular differentiation and tissue regeneration. In comparison to materials with extra-fibrillar mineralization, those with intra-fibrillar mineralization demonstrate enhanced effects in inducing osteogenic differentiation of stem cells and promoting bone regeneration^[21, 22]^. To achieve enhanced tissue regeneration effects, *in vitro* bio-inspired mineralization techniques have been gradually developed. By adjusting the collagen molecule solution from acidic to alkaline conditions, collagen fibrils with a periodic d-space structure of 67 nm can be obtained, offering a template for achieving bio-inspired mineralization^[9, 10]^. Polyelectrolytes, such as polyaspartic acid (pAsp) and polyacrylic acid, plays a similar role to natural NCPs and has been widely^[10]^. Once stabilized by polyelectrolytes, the ACP can enter the ≈ 1.8 – 4 nm sized tortuous subchannels of the fibrils^[10, 23]^. Subsequently, ACP undergoes transformation into crystallized CaP, gradually forming a hierarchical and oriented structure. Finally, the ultrastructural features of the mineralized constructs reveal the accumulation of minerals in both intra- and extrafibrillar compartments, resembling bone. The highly mineralized scaffold can offer an optimized microenvironment for osteogenic induction^[21, 24]^ and bone growth^[21]^. However, it remains unclear the effect of different mineralized collagen scaffold on inducing the differentiation of stem/ progenitor cells into fibrochondrocytes and the regeneration of fibrocartilage tissue.

Here we utilized FIB-SEM to decode the nanostructure of the TBI. Subsequently, through the development of a “floating mineralization” system, we created a bio-inspired mineralization collagen hydrogel (BIMCH) consisting of three layers that replicate the hierarchical nano- to micro-scale architecture of TBI. The middle layer, comprising approximately 34 wt% of inorganic content, is particularly noteworthy for its resemblance to the nanostructure of fibrocartilage and its exceptional capacity to induce mineralized fibrochondrogenesis *in vitro*. Through motor function analysis, imaging diagnosis, histological staining, immunofluorescence staining, and biomechanical assessments, we provide evidence that the in situ transplantation of the novel hydrogel achieves synchronized tendon-fibrocartilage-bone regeneration, resulting in a remarkable 68% maximum mechanical recovery at 8 weeks post-operation. Single-cell RNA sequencing analysis reveals that a unique atlas of in situ stem/progenitor cells is generated during the TBI healing *in vivo*. Notably, by single-cell RNA analysis, we show that the BIMCH with an ability to direct endogenous cells, Gli1^+^Dkk3^+^progenitor cells, to facilitate TBI regeneration (Fig.1).

**Fig. 1.**
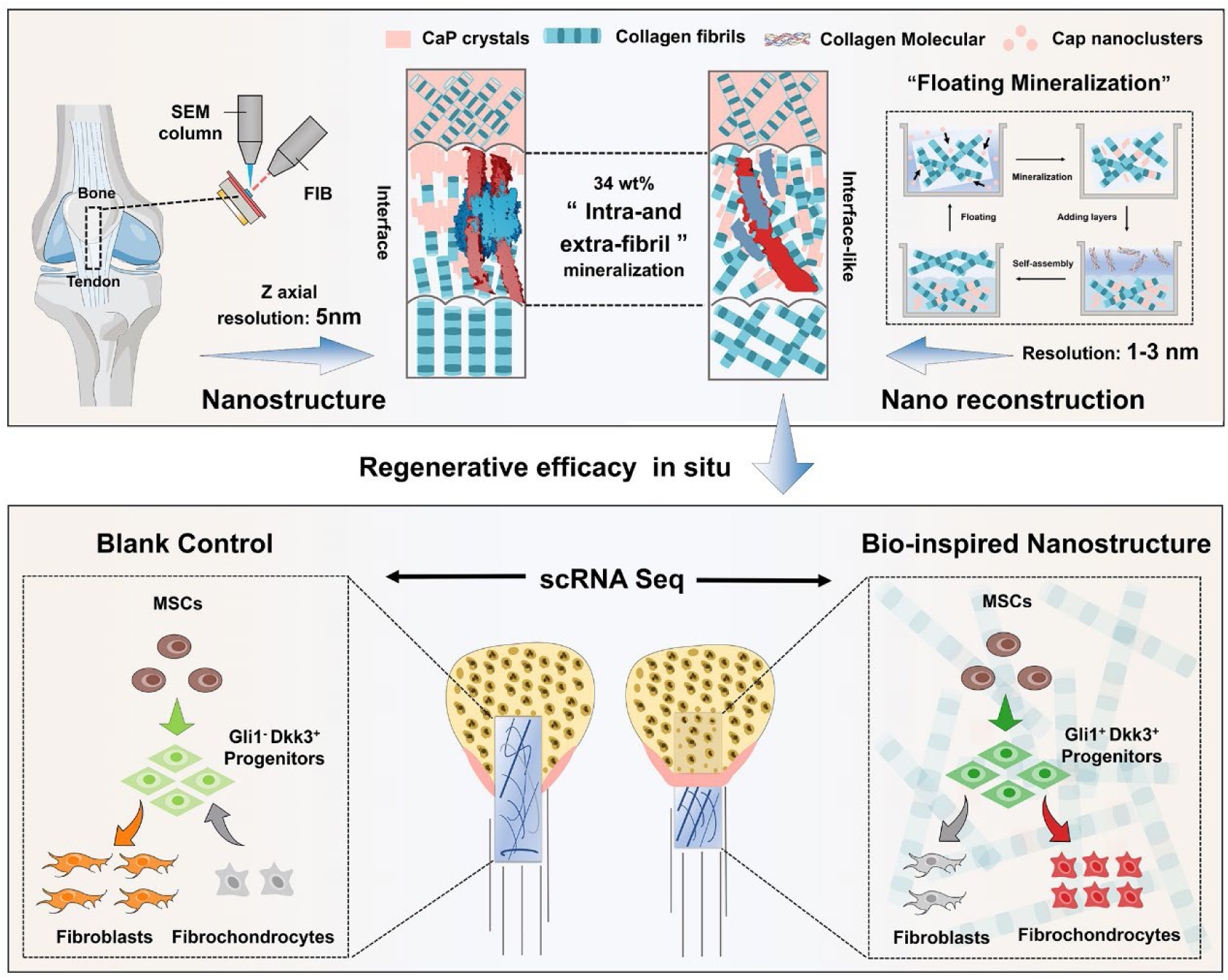
An overview of the vision. The three-layer hydrogel, which mimics the hierarchical nano- to micro-scale structure of TBI. Following in situ implantation, the hydrogel was capable of guiding endogenous cells, specifically Gli1+Dkk3+ progenitor cells, to promote TBI regeneration.

## Results

### The compositional and 3D nanostructure of the tendon–bone interface

To acquire precise biophysical and biochemical cues for constructing biomimetic scaffold, we identified the nanoscale compositions and assembly patterns of the enthesis. As expected, the three regions displayed varying contents of calcium (Ca) and phosphate (P) elements (Fig.2 a, b). The Ca and P gradually decreased from the bone area to the tendon area (Fig.2 a, b). Subsequently, 2D (Fig.2 d) and 3D (Fig.2 f-h) projections of the apatite mineral phase were analyzed. Interestingly, we identified three distinct motifs of mineral organization (Fig.2 d, f): the complete mineralization of collagen fibrils in the bone (Supplementary Fig.1), the partial mineralization of collagen fibrils in the interface, the collagen fibrils without mineral in the tendon. To explore the organization and relationship between the principal components of mineralized tissues, namely mineral and collagen, we selected specific regions of interest for 3D reconstruction. Whether complete or partial mineralization of collagen fibrils, mineral particles do not exclusively align intrafibrillarly or extrafibrillarly; instead, they form a continuous cross-fibrillar phase (Fig.2 g(i-ii), Fig.2 h(i-ii), Supplementary Fig.2). Furthermore, the mineral appeared as irregularly elongated shape assemblies (Fig.2g(iii), Fig.2 h(iii)).

**Fig. 2.**
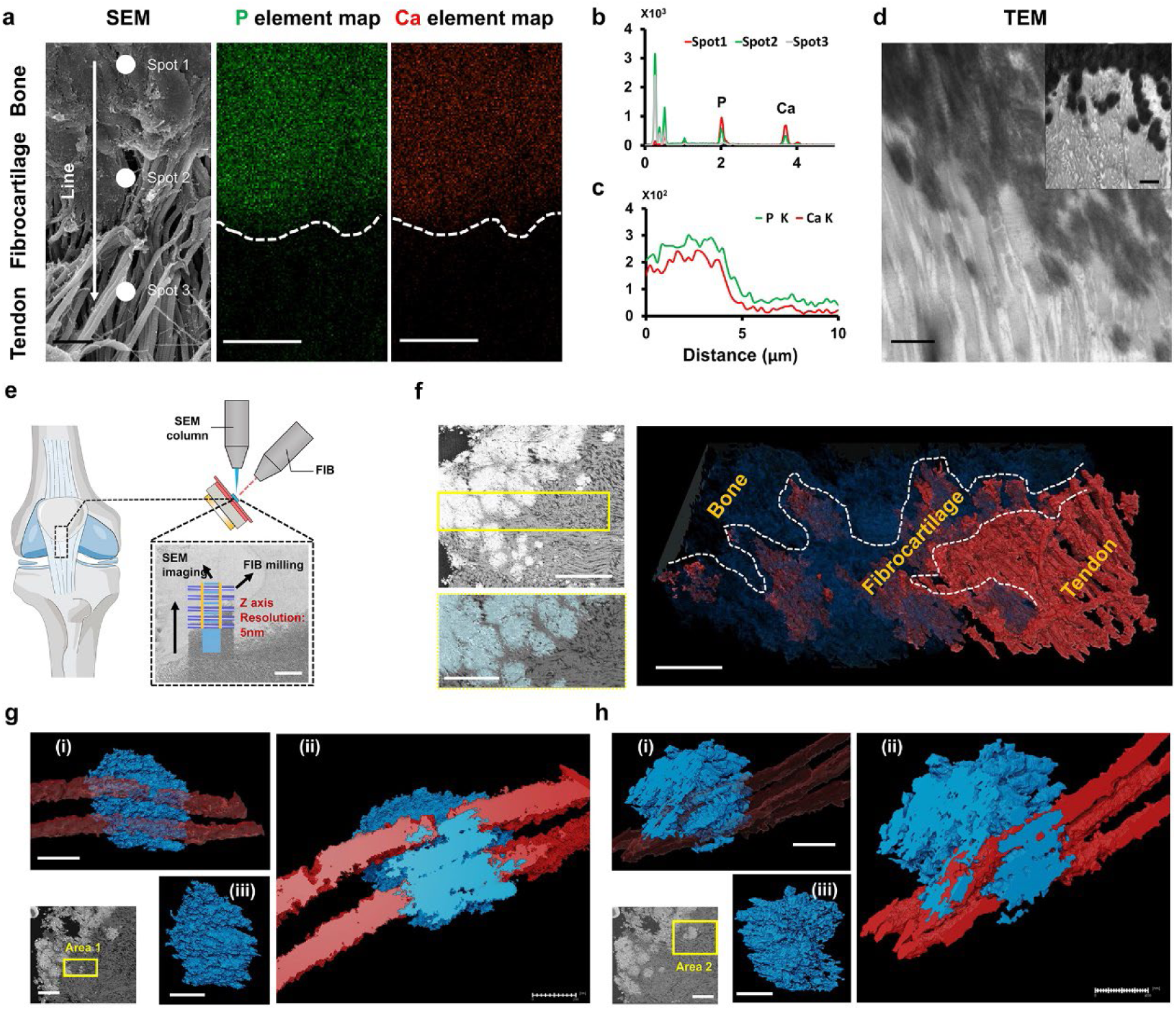
The high-resolution and three-dimensional images of the enthesis. **a-c,** SEM image and EDX mappings of elemental distribution were recorded (**a**). The EDX spectral confirmed a gradient in elemental content within each spot (**b**) and along the vertical axis from the bone to the tendon (**c**), scale bar: 1 μm. **d,** TEM images of the TBI reveal distinct degrees of mineralization, with dark regions indicating areas of mineral deposition. scale bar: 500 nm. **e,** The focused ion beam scanning electron microscopy (FIB-SEM) technique employs a FIB to first expose a surface within the enthesis, followed by imaging using SEM. This iterative process is repeated thousands of times. **f**, The left panel displays the selected area (yellow box) with a scale bar of 3 μm, while the right panel showcases corresponding 3D reconstructions in the TBI with a scale bar of 2 μm. Collagen fibrils are depicted in red, and mineralization is represented in blue. **g**, The morphology and organization of crystals within the fibrocartilage are displayed as translucent fibrils (**i**, scale bar: 300 nm) or hemi-sectioned fibrils (**ii**, scale bar: 200 nm). In (**iii**), only the mineral deposits are displayed with a scale bar of 300 nm. **h**, The morphology and organization of crystals within the fibrocartilage are illustrated as translucent fibrils (**i**, scale bar: 600 nm) or hemi-sectioned fibrils (**ii**, scale bar: 400 nm). In (**iii**), only the mineral deposits are displayed with a scale bar of 500 nm.

### Bio-inspired tendon–bone interface with gradient nanostructure

To create collagen hydrogels, we prepared gels using acid-soluble type I collagen extracted from pig tendons, employing ammonia vapor diffusion in cylindrical molds (Fig. 3a). To induce mineralization in these collagen gels, they were immersed in a solution containing Ca, P, and pAsp (Fig. 3a). To quantify the mineral content, thermogravimetric analysis (TGA) was employed. Over time, the mineral content of the gels increased progressively, from 5.48 ± 2.71 wt % to 24.17 ± 2.01 wt %, 34.19 ± 2.52 wt %, 45.72 ± 0.97 wt %, and finally to 61.04 ± 0.77 wt % (Fig. 3b, Supplementary Fig.3a). Subsequently, to mimic the three distinct mineral organization patterns found in native TBI, we selected collagen gels mineralized for 0 days, 2 days, and 6 days as the tendon-like (TL) layer, fibrocartilage-like (FCL) layer, and bone-like (BL) layer, respectively. The BL layer exhibited an elastic modulus of 8.42 ± 7.9 KPa, while the FCL layer demonstrated an elastic modulus of 43.17 ± 68.15 KPa. In contrast, the BL layer displayed a substantially higher elastic modulus of 120.00 ± 83.52 KPa (Fig. 3c). Our “floating mineralization system” enabled the creation of multi-layered scaffolds, offering precise control over the spatial distribution of mineralized collagen, as well as the thickness and number of layers. After gelation, fixation, and mineralization, additional layers could be added by acid-soluble solution collagen on top of the existing gel (Fig. 3a). This resulted in a continuous, interlocked hydrogel comprising three layers: the TL layer with minimal Ca, the FCL layer with moderate Ca content, and the BL layer with a significant Ca presence (Fig. 3d, Supplementary Fig.3b and c).

**Fig. 3.**
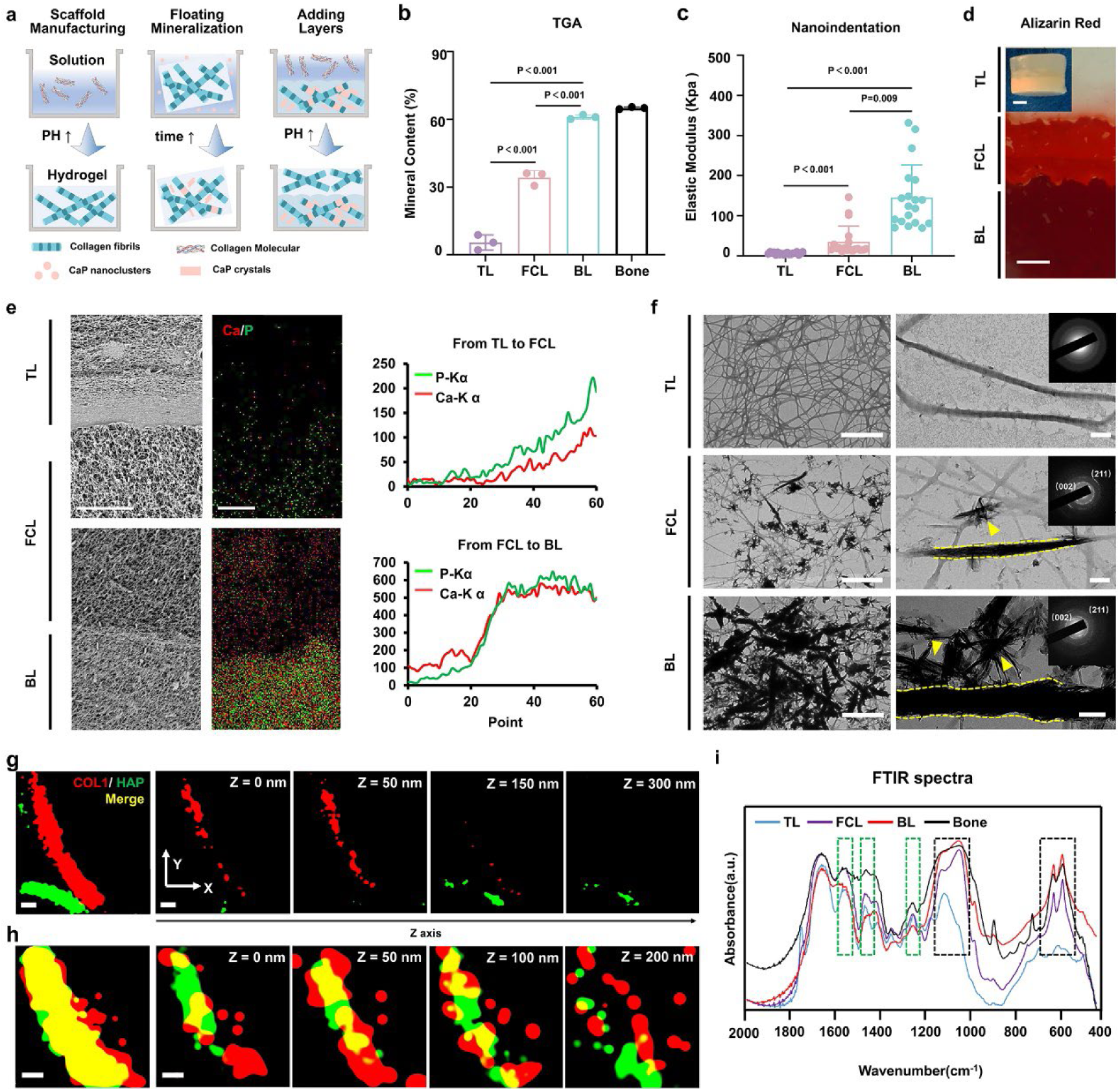
The bio-inspired tendon–bone interface with gradient nanostructure. **a,** Schematic illustrating the fabrication process of the novel collagen hydrogel. **b**, Thermogravimetric analysis of the three layers and native bone (n = 3). **c**, Nanoindentation modulus of the three layers (n = 19). **d**, Alizarin Red staining images of the gel demonstrate continuous, interlocked and three-layer structure. Scale bar: 1.5 mm. **e**, SEM micrographs (left panel, scale bar: 10 μm), EDX mapping of elemental distributions (middle panel, scale bar: 5 μm), and elemental content (right panel) were recorded from the cross-section of the graded mineral scaffold. **f**, TEM showed bio-inspired mineralization, with intrafibrillar mineralization and extrafibrillar mineralization indicated by the yellow dashed line and arrowheads, respectively. Selected area diffraction reveals oriented crystals of hydroxyapatite with the c-axis aligned with the long axis of the collagen fibril (inset of the right panel). Left panel, scale bar: 2 μm; right panel, scale bar: 200 nm. **g**, The 2D and z-slice of the STORM images illustrate the extrafibrillar mineralization. scale bar: 100 nm. **h**, The 2D and z-slice of the STORM images illustrate the intrafibrillar mineralization. scale bar: 100 nm. **i**, FTIR spectra of the of the three layers compared to native bone. The green dashed line represented the peak position of amide I, II, and III (1649, 1553, and 1248 cm^−1^) bands, while the dark dashed line represented the peak position of apatitic phosphate (1030, 600, and 560 cm^−1^) bands.

Scanning electron microscopy (SEM) analyses revealed that both of the three layers exhibited a fibrillar and sponge-like structure with a high degree of porosity and interconnectivity, featuring pore sizes ranging from 1 to 3 µm (Supplementary Fig.3c, left panel). With increased mineralization time, there was a greater presence of distributed extrafibrillar Ca and P deposits, resulting in a rougher structure (Supplementary Fig.3c, middle and right panel). Subsequently, we characterized the interface of scaffold cross-sections, which displayed a visible distinction among the three layers (Fig. 3e). Notably, even at the microscale, there was no distinct gap between adjacent layers, indicating strong mechanical interlocking (Fig. 3e). Furthermore, both Ca and P content exhibited a gradual increase from the TL layer through the FCL layer to the BL layer (Fig. 3e).

The TL layer, with fibril diameters measuring 60.6 ± 12.0 nm (n = 50), exhibited the typical 67 nm D-period (Fig. 3f). In contrast, the diameter of mineralized fibrils in the FCL layer measured 117.0 ± 40.1 nm (n = 50), while in the BL layer, it measured 171.5 ± 49.2 nm (n = 50) (Fig. 3f). Similar to the native TBI, there were three distinct motifs of mineral organization (Fig. 3f): collagen fibrils without mineral in the TL layer, partial mineralization of collagen fibrils in the FCL layer, and complete mineralization of collagen fibrils in the BL layer. Furthermore, in both the FCL and BL layers, we observed both intra- and extrafibrillar localization of apatite crystallites’ (Fig. 3f). Selective area electron diffraction (SAED) analyses of mineralized fibrils also revealed the typical broad arcs for the (002) and (211) planes, consistent with the known hexagonal crystal form of hydroxyapatite found in native bone and osteoblast-secreted apatite crystals^[11]^. The 3D-HR images can be obtained by using stochastic optical reconstruction microscopy (STORM), allowing for the identification of HAP minerals both on the external (Fig. 3g) and internal (Fig. 3 h) portion of the collagen-I fibrils. In the last, we characterized the chemical composition of our hydrogel in comparison to that of native bone using fourier transform infrared spectroscopy (FTIR). All three layers exhibited characteristic peaks corresponding to amide I, II, and III (1649, 1553, and 1248 cm^−1^) bands, exhibited characteristic peaks corresponding to amide I, II, and II. Additionally, the FCL and BL layers displayed characteristic peaks corresponding to apatitic phosphate (1030, 600, and 560 cm^−1^) bands (Fig. 3i).

In summary, a gradient bio-inspired mineralization collagen hydrogel (BIMCH) was developed. This gel incorporates three distinct motifs of mineral organization: collagen fibrils devoid of minerals in the TL layer, partially mineralized collagen fibrils in the FCL, and fully mineralized collagen fibrils in the BL layer. At the micro-scale, the hydrogel exhibited a range of gradient properties, including mineral content (ranging from 5% to 34% to 61%), mechanical stiffness (ranging from 8 KPa to 43 KPa to 119 KPa), and surface roughness (ranging from smooth to rough to rougher). At the nano-scale, fibrils without minerals exhibited the typical 67 nm D-period, whereas mineralized fibrils featured intra- and extrafibrillar minerals.

### The seeded stem cells underwent differentiation into tenocytes, mineralized fibrochondrocytes, and osteoblasts following cultivation in the TL, FCL, and BL layers, respectively

To investigate the biological effects of the BIMCH, mesenchymal stem cells (MSCs) were seeded on TL layer, FCL layer, and BL layer, respectively (Fig. 4a). Live/dead staining results clearly indicated that approximately 90% of cells cultured on these three layers remained viable even after at least 7 days of *in vitro* culture, demonstrating the favorable biocompatibility of the BIMCH (Fig. 4b, c). Subsequently, after 3 days, we conducted RNA-seq analysis to comprehensively and impartially examine the early events associated with MSCs’ response to bio-inspired mineralization. The analysis of differentially expressed genes (DEGs) revealed that the three layers had a significant impact on the gene expression of MSCs (Supplementary Fig.4a). Furthermore, hierarchical clustering and principal component analysis (PCA) demonstrated that cells of the same type from the three groups clustered together based on their distinct gene expression profiles (Fig. 4d). DEGs were further categorized based on their responses to varying levels of bio-inspired mineralization and the direction of that response (Supplementary Fig.4b). We identified ten different types of expression change patterns among DEGs across the three groups, ranging from down-down profiles to up-up profiles, down-up profiles, up-down profiles, down profiles (FCL vs. BL), up profiles (FCL vs. BL), down profiles (TL vs. FCL), up profiles (TL vs. FCL), down profiles (TL vs. BL), and up profiles (TL vs. BL). In comparison to the TL group, a greater number of genes were found to be upregulated or downregulated in the FCL or BL groups, which was consistent with the Venn Diagram (Supplementary Fig.4c). Noteworthy genes in the “Up profile” (TL vs. BL) included well-known proliferative genes like *Mcm4*, while the “Down profile” (TL vs. BL) featured genes associated with cellular stress, such as *Hspa5*, *Hsp90b1*, and *Hyou1*.

**Fig. 4.**
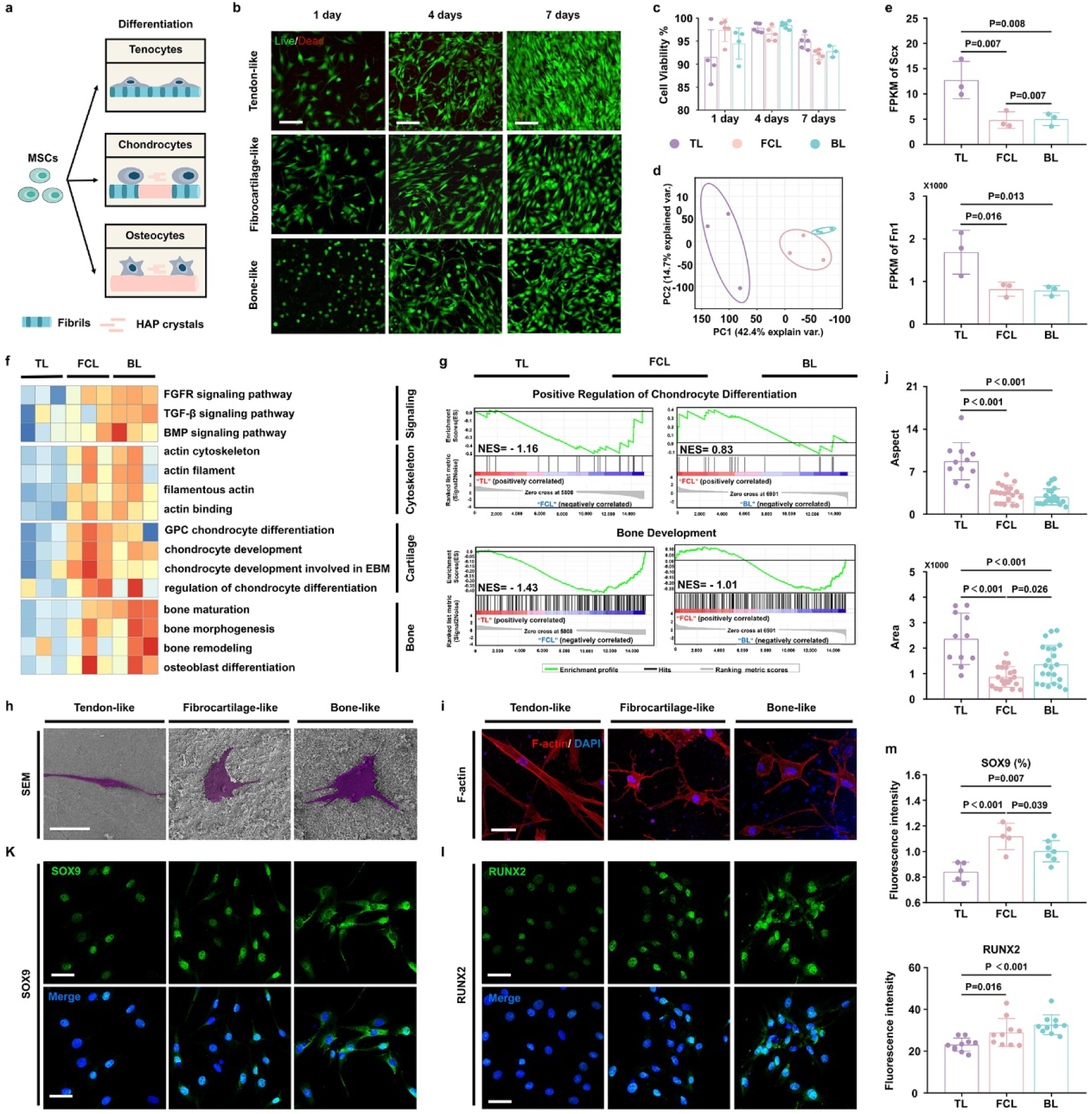
BIMCH induced MSCs toward TBI cell lineage differentiation *in vitro*. **a**, Schematic overview of *in vitro* cell differentiation analysis. **b**-**c,** Representative fluorescence images depicting live-dead MSCs (**b**), quantitative analysis (**c**) and after culturing in TL layer, FCL layer, and BL layer for 1, 4, and 7 days. after culturing in TL layer, FCL layer, and BL layer for 1, 4, and 7 days. Green and red indicate live and dead cells, respectively. Scale bar, 100 μm. (n = 6). **d**, PCA of all samples using gene expression revealed three distinct expression patterns. **e**, The FPKM values of tendon-related genes in the three groups. **f**, Heatmap displaying gene sets related to signaling pathways, cytoskeleton, cartilage, and bone development, based on Gene Set Variation Analysis (GSVA) enrichment analysis. **g**, Gene Set Enrichment Analysis (GSEA) of genes associated with chondrocyte differentiation and bone development. **h-j**, Representative SEM (**h**) and fluorescence images (**i**) of MSCs morphology after 1 day of culturing on the TL, FCL, and BL layers, respectively. Cell morphology quantified for cell area and cell aspect (**j**) in each group (n = 3). Scale bar for the fluorescence images: 40 μm; Scale bar for the SEM images: 30 μm. **k**-**m**, Representative SOX9 staining of MSCs seeded on the TL (tendon-like), FCL, and BL (bone-like) layers after 7 days of culture.

Next, we conducted a more in-depth analysis to gain functional insights into the behavior of MSCs after 3 days of culturing on the three distinct layers. Among the three groups, the tendon-related genes, such as *Scx* and *Fn1*, were significantly highest in the TL group (Fig. 4e). In contrast, when compared with the TL group, the biological processes associated with signaling pathways, cytoskeleton, cartilage, and bone were significantly more enriched in the FCL and BL groups (Fig. 4f, g). Notably, among these three groups, the biological processes related to cartilage and bone were most significantly enriched in the FCL and BL groups, respectively. MSCs on the TL scaffold exhibited a spindle-shaped “tenoblast-like” morphology with long tapering cytoplasmic extensions, while cells on the BL scaffold displayed a branched “osteocyte-like” shape with long filopodia (Fig. 4h-j). In contrast, cells on the FCL scaffold exhibited distinct shapes, and there were differences in cell area and cell aspect compared to the “tenoblast-like” and “osteocyte-like” shapes. After 7 days of culturing on the scaffolds, MSCs were subjected to immunostaining for SOX9 and RUNX2 (Fig. 4k-m). Following quantitative analysis of fluorescence intensity, it became evident that both SOX9 and RUNX2 levels were significantly increased in the FCL and BL groups when compared to the TL group. Among the three groups, the expression of SOX9 was significantly highest in the FCL group, while the expression of RUNX2 was significantly highest in the BL group.

In summary, when cultured in the TL, FCL, and BL layers, the seeded stem cells underwent differentiation into tenocytes, mineralized fibrochondrocytes, and osteoblasts, respectively.

### The transplantation of the bio-inspired mineralization collagen hydrogel promotes the synchronous regeneration of the tendon-fibrocartilage-bone complex

We translated both the non-mineralized collagen hydrogel (NMCH) and BIMCH into an *in vivo* rat model simulating the patellar-to-tendon interface fenestration (PTF) to assess the regeneration of the TBI (Fig. 5a and Supplementary Fig5). One week post-surgery, the BIMCH group exhibited higher Achilles Functional Index (AFI) and grid-hanging time values compared to the blank control (BC) group, indicating excellent repair performance (Supplementary Fig6). However, by the end of the second week post-surgery, there were no significant differences in AFI and grid-hanging time values among the three groups (Supplementary Fig6).

**Fig. 5.**
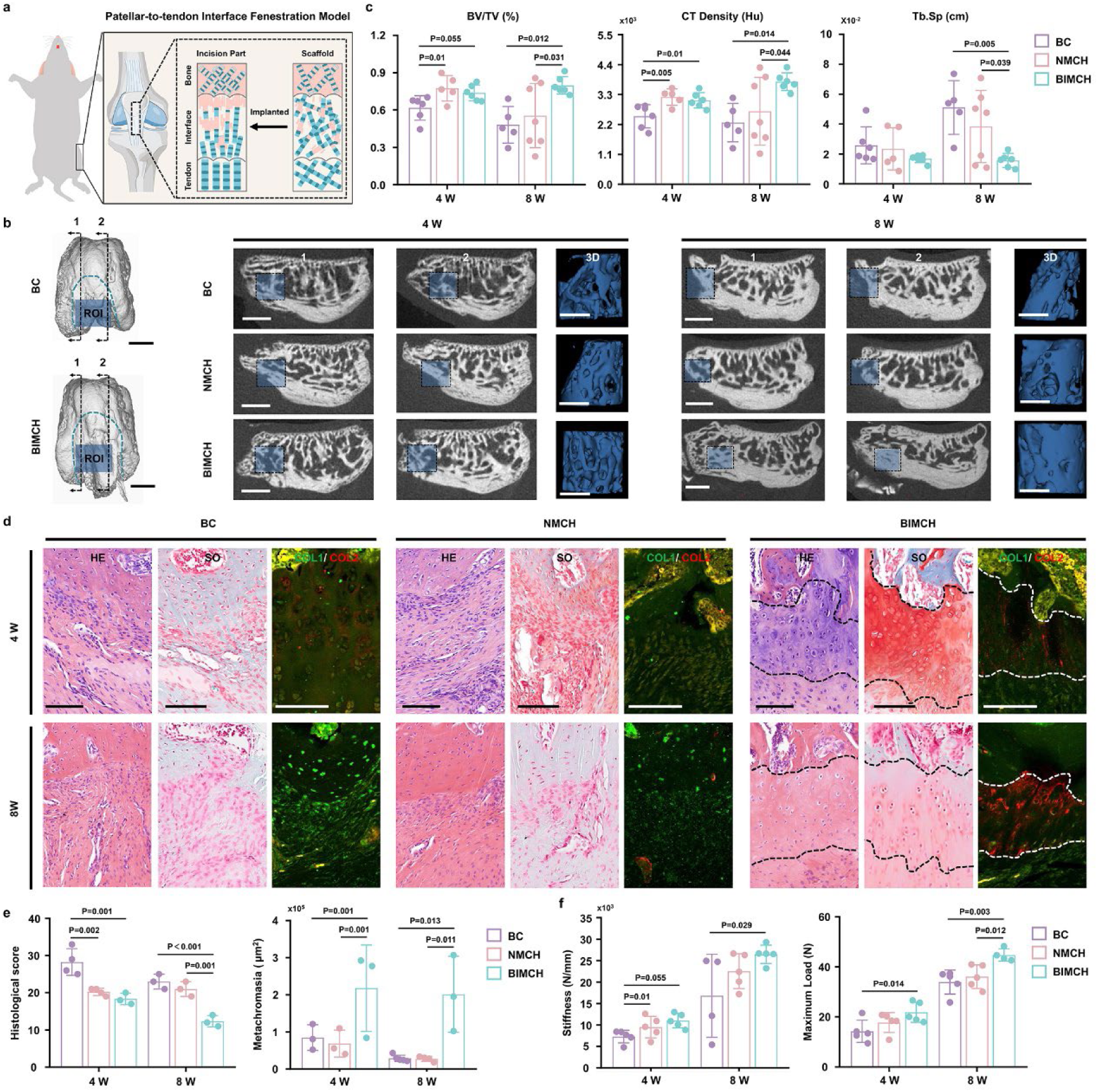
BIMCH enhances regeneration of rat tendon-bone interface defect. **a,** Schematic overview of TBI regeneration efficacy *in vivo*. **b,** At 4 and 8 weeks post-operation, sagittal-section and 3D reconstruction micro-CT images of the specified region (highlighted in blue with dashed lines) for the BC, NMCH, and BIMCH groups. Scale bars: 1 mm. **c,** Micro-CT quantification of the specified region, including BV/TV and Tb.Sp. for BC, NMCH, and BIMCH samples (n = 5-7). **d**, Representative images of sections stained with H&E (left panel), SO (middle panel), COL1/COL2 (right panel) from rats treated with BC, NMCH, and BIMCH. **e**, Assessment of histological maturity and analysis of the area of newly formed fibrocartilage in HE and SO staining images from the three groups (n = 3-4). **f**, Stiffness, maximum load values of the repaired interface tissues (n = 4-5).

After euthanizing the rats at 2, 4, and 8 weeks post-surgery, no discernible signs of ectopic ossification or infection were observed in the implantation area (Supplementary Fig7,8). Examination of patella sagittal-section and 3D reconstruction images of the region of interest revealed a greater amount of new bone formation in both the NMCH and BIMCH groups compared to the BC group at 4 weeks post-operation by micro-CT scanning (Fig. 5b). Quantitative analysis demonstrated that the CT density in the NMCH and BIMCH groups was significantly higher than that in the BC group (Fig. 5c). By 8 weeks post-operation, abundant newly formed bone had regenerated in the BIMCH group (Fig. 5b). In comparison to the other two groups, the bone volume over total volume (BV/TV), CT density, and trabecular spacing (Tb.Sp.) values of the BIMCH-implanted group exhibited significant increases (Fig. 5c).

By week 4, newly formed collagen bundles in the TBI appeared no significant difference between the three groups (Supplementary Fig9). By week 8, the collagen fibers exhibited greater organization in the BIMCH group (Supplementary Fig9). The gray values of the BIMCH-implanted group were significantly higher compared to the BC and NMCH groups. Secondly, tissue morphology was assessed using H&E and SO stainings. The repaired BTI remained defective in the BC group but was intact in the NMCH and BIMCH groups (Supplementary Fig10). In the NMCH group, noticeable fibrovascular granulation tissue and inflammatory cellular infiltration were observed at the insertion site (Fig. 5d, Supplementary Fig10). In the BIMCH specimens, fibrocartilaginous tissue was evident, although the chondrocytes appeared mostly immature and disorganized compared to the sham group (Fig. 5d, Supplementary Fig10). The total histologic score of the BC group was significantly higher than that of the NMCH and BIMCH groups (Fig. 5e). By week 8, the chondrocytes appeared more mature and organized compared to week 4 (Fig. 5d). The total histologic scores of the BC and NMCH groups were significantly higher than that of the BIMCH groups (Fig. 5e). Thirdly, to further elucidate the presence of newly formed fibrocartilaginous tissue, all samples underwent SO staining. The area of metachromasia at the repair site significantly increased in the BIMCH group compared to the BC and NMCH groups, indicating improved new fibrocartilage formation (Fig. 5d, e). Lastly, the BIMCH group exhibited region-dependent levels of collagen type I and II at the tendon–bone insertion at 4 and 8 weeks post-operation (Fig. 5d, Supplementary Fig11). The fluorescence intensity plot over distance showed that the collagen type I-lacking region was complementary to the collagen type II-rich region in BIMCH group, which is as similar as native TBI tissue (Fig. 5d, Supplementary Fig11).

The biomechanical experiments have confirmed the healing of the TBI in the BIMCH group (Fig. 5f). Tension and toughness exhibited an increase over time, with values being notably greater at 8 weeks compared to 4 weeks. Furthermore, both the maximum load and stiffness values of the BIMCH group were significantly higher than those of the BC group at both week 4 and week 8. By week 8, the maximum load values of the BIMCH group were also significantly higher than those of the NMCH group, and result in 68% recovery of the native group (65.62 ± 8.79 N, n = 4).

In summary, BIMCH induced significant collagen alignment, fibrocartilage and bone ingrowth, thereby exhibiting exceptional mechanical properties for load-bearing tendon-to-bone connections.

### Single-cell transcriptomes of the enthesis regenerative microenvironment revealed the presence of four distinct cell clusters

To elucidate differentially regulated cell subsets in response to BIMCH treatment, we conducted scRNA-seq analysis on the in situ cell clusters after a 2-week implantation of the cell-induced differentiation system (Fig. 6a). Following quality control and doublet exclusion, we obtained data from 15,866 cells: 9,619 cells in the BC group and 6,247 cells in the BIMCH-treated group.

**Fig. 6.**
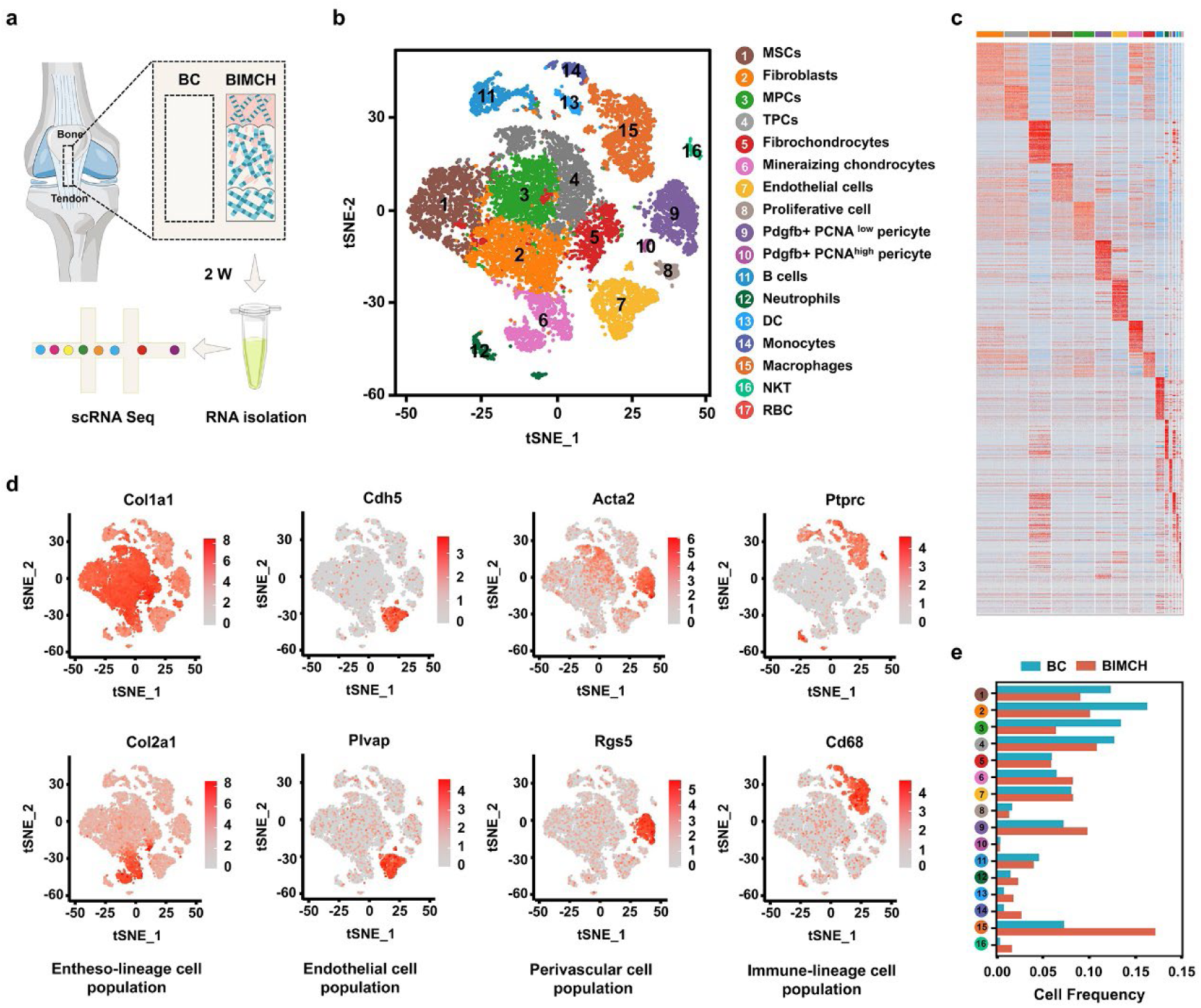
A single-cell atlas of the TBI regenerative microenvironment. **a,** Workflow illustrating the isolation of samples and single-cell RNA sequencing of regenerated tissue at 2 weeks post-defect surgery. n = 4 samples per group. **b,** tSNE plot shows the distribution of 15866 cells that divided into 17 sub-clusters. MSCs: mesenchymal stem cells; MPCs: mesenchymal progenitor cells; TPCs: tendon progenitor cells. **c,** Heatmap shows expression of marker genes of the 17 sub-clusters. Marker genes were selected according to p value (Wilcoxon Rank Sum test, top 50). **d,** tSNE plots of the expression of specific genes, which mark four different cell populations. **e,** Barplots of the relative cell frequency of 17 sub-clusters between the BC and BIMCH groups, respectively.

Unsupervised clustering identified 17 distinct clusters (Fig. 6b). These subpopulations were categorized into four groups: an entheso-lineage cell population (expressing *Cola1* and *Col2a1*); an endothelial cell population (expressing *Cdh5* and *Plvap*); a perivascular cell population (expressing *Acta2* and *Rgs5*); and an immune-lineage cell population (expressing *Ptprc* and *Cd68*) (Fig. 6d). Subsequently, we examined the differentially regulated cell subpopulations following in situ implantation of BIMCH. A significant difference in the proportion of immune-lineage and entheso-lineage cells was observed between the BC and BIMCH groups (Fig. 6e).

### Elucidating entheso-lineage specification during TBI regeneration

The cellular composition and differential regulation of entheso-lineage cell subpopulations were influenced by BIMCH (Fig. 7 a). Within the entheso-lineage cells, we identified eight major subsets through sub-clustering analysis (Fig. 7 b). In comparison to the BC group, the BIMCH group exhibited a significant increase in the proportion of clusters 6-8, along with a decrease in clusters 4-5 subpopulations (Fig. 7 c). We then identified two Nov+ mesenchymal stem cells (MSCs) populations (clusters 1-2), one of which exhibited a high level of *Thy1* (Thy1^+^ MSCs, cluster 2) (Fig. 7 d, g, h). Gene ontology (GO) analysis revealed that MSCs were enriched with genes regulating metabolic processes, whereas Thy1^+^ MSCs were associated with cell migration (Fig. 7e). Tendon progenitor cells (TPCs) were defined based on their expression of *Thy1*, *Scx*, *Tnmd*, and *Postn* (Fig. 7 d, g). Cluster 5 represented a mesenchymal subset expressing *Dkk3* and *Cxcl14* (Fig. 7 d, g, h), and had the highest StemID score among the eight clusters, consequently classified as mesenchymal progenitor cells (MPCs). GO analysis indicated that MPCs were enriched with genes regulating skeletal muscle tissue regeneration (Fig. 7e).

**Fig. 7.**
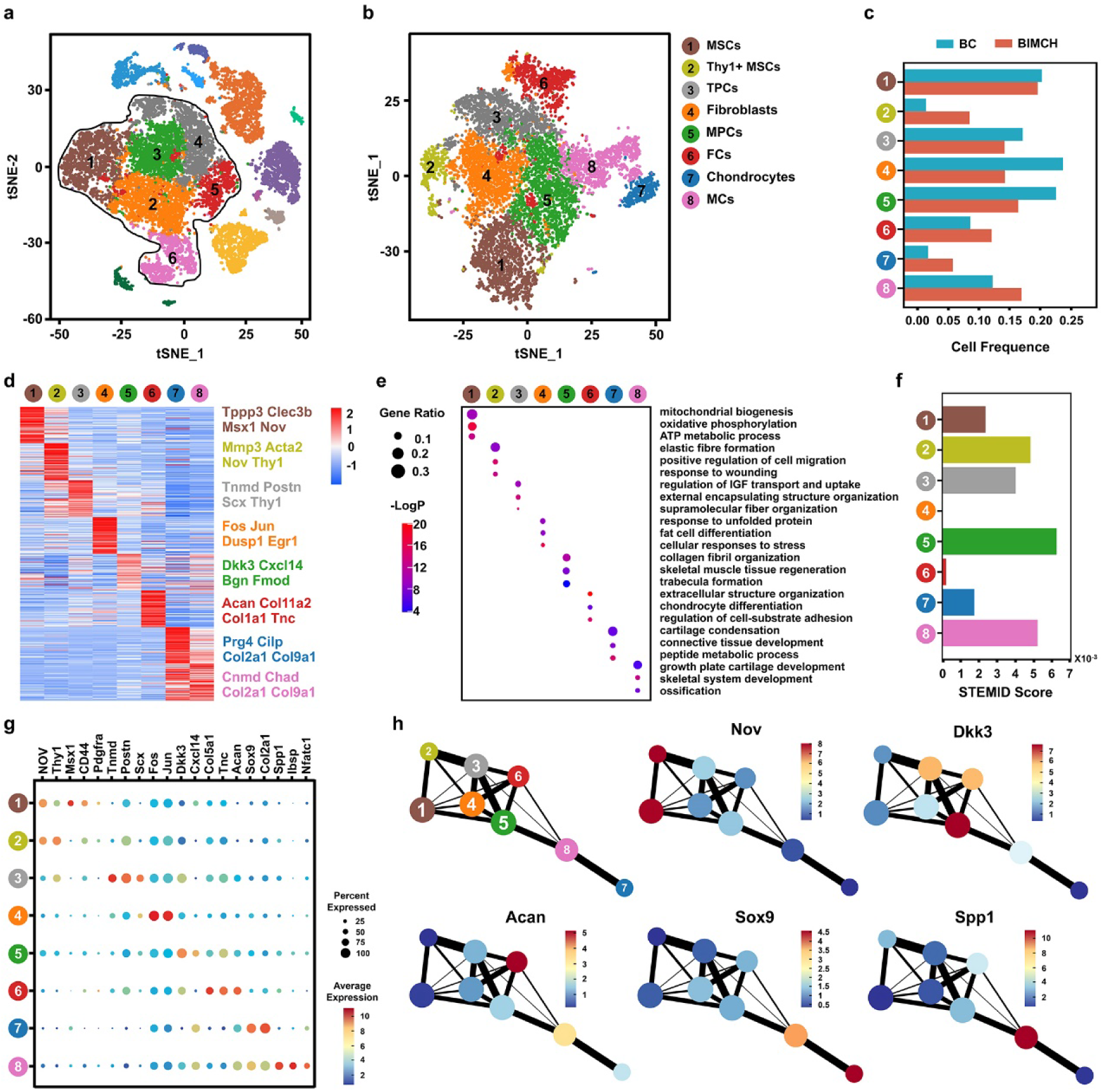
Characterization of the entheso-lineage. **a,** t-SNE plot depicting the distribution of 15,866 cells, with the entheso-lineage cells highlighted (dark dotted line). **b,** Visualization of entheso-lineage cells highlighted in (a) using a t-SNE plot, further divided into eight sub-clusters. MSCs: mesenchymal stem cells), MPCs: mesenchymal progenitor cells, TPCs: tendon progenitor cells, FCs: fibrochondrocytes, MCs: mineralized chondrocytes. **c,** Barplots displaying the relative cell frequency of the eight sub-clusters between the BC and BIMCH groups. **d**, Heatmap illustrating the expression of marker genes for the eight sub-clusters. Marker genes were selected based on p-values (Wilcoxon Rank Sum test, top 50). **e,** Enriched Gene Ontology (GO) terms for differentially expressed genes (DEGs) in the entheso-lineage clusters. **f**, Barplots depicting StemID scores for entheso-lineage clusters. **g**, Dot plots displaying the expression of curated feature genes in the eight subsets. Dot size represents the proportion of cells expressing a specific gene in the indicated subset, and the color bar indicates gene expression levels. **h**, Partition-based graph abstraction (PAGA) illustrating the connectivity among subsets in (b). The mean expression of representative genes (Mesenchymal: *Nov*; Chondrogenic: *Acan* and *Sox9*; Osteogenic: *Spp1*) in each subset is shown in the abstracted graph. Line thickness indicates the strength of connectivity, and the color bar represents gene expression levels.

Fibroblasts (FCs, cluster 4) were identified based on their expression of *Fos* and *Jun*. Cluster 6 co-expressed tenogenic (e.g., *Col1a1*, *Tn1*) and chondrogenic markers (e.g., *Acan*) (Fig. 7 d, g, h) and exhibited enriched profiles of chondrocyte differentiation (Fig. 7e), thus being defined as fibrochondrocytes (FCs). We also identified signature profiles of chondrocytes (e.g., *Sox9*, *Col2a1*, cluster 7) and mineralizing chondrocytes (e.g., *Sox9*, *Col2a1*, *Spp1*, *Ibsp*, *Nfatc1*, MCs, cluster 8). GO analysis revealed that chondrocytes were enriched with genes regulating cartilage condensation, while MCs were associated with growth plate cartilage development (Fig. 7e). Partition-based graph abstraction (PAGA) analysis indicated the pivotal role of MPCs (Dkk3+) in connecting Nov+ MSCs to FCs, chondrocytes, or MCs (Fig. 7h).

### Gli1+Dkk3+ progenitor cells activated by bio-inspired mineralization nanostructure for TBI regeneration

Initially, pseudotime analysis using RNA velocity was employed to investigate lineage relationships among the entheso-lineage cell subsets. In the natural healing condition without treatment, MPCs differentiated from other cell subpopulations and were more abundant than in the BIMCH-treated group (Fig. 8a). While, in the BIMCH healing condition, strong directional streams were observed from MSCs towards MPCs, eventually leading to fibroblasts, MCs, and FCs, forming three differentiation trajectories (Fig. 8a). Diffusion map analysis of entheso-lineage cell subsets revealed two differentiation trajectories involving fibrogenesis and chondrogenesis (Fig. 8b). When MSCs were designated as the root to identify genes temporally expressed over pseudotime, it was observed that genes highly expressed in MSCs (e.g., *Nov*, *Thy1*, *Msx1*) gradually down-regulated, while genes highly expressed in fibrogenesis (e.g., *Dkk3*, *Postn*, *Tnmd*, *Scx*, and *Thbs4*) and chondrocytes (e.g., *Acan*, *Sox9*, *Col2a1*, *Col9a1*, and *Spp1*) were upregulated during terminal differentiation (Fig. 8c). Notably, *Dkk3* and *Postn* were upregulated during the early-middle differentiation process leading to fibrogenesis, whereas *Col2a1* and *Sox9* were upregulated during the later differentiation process leading to chondrocytes (Fig. 8d).

**Fig. 8.**
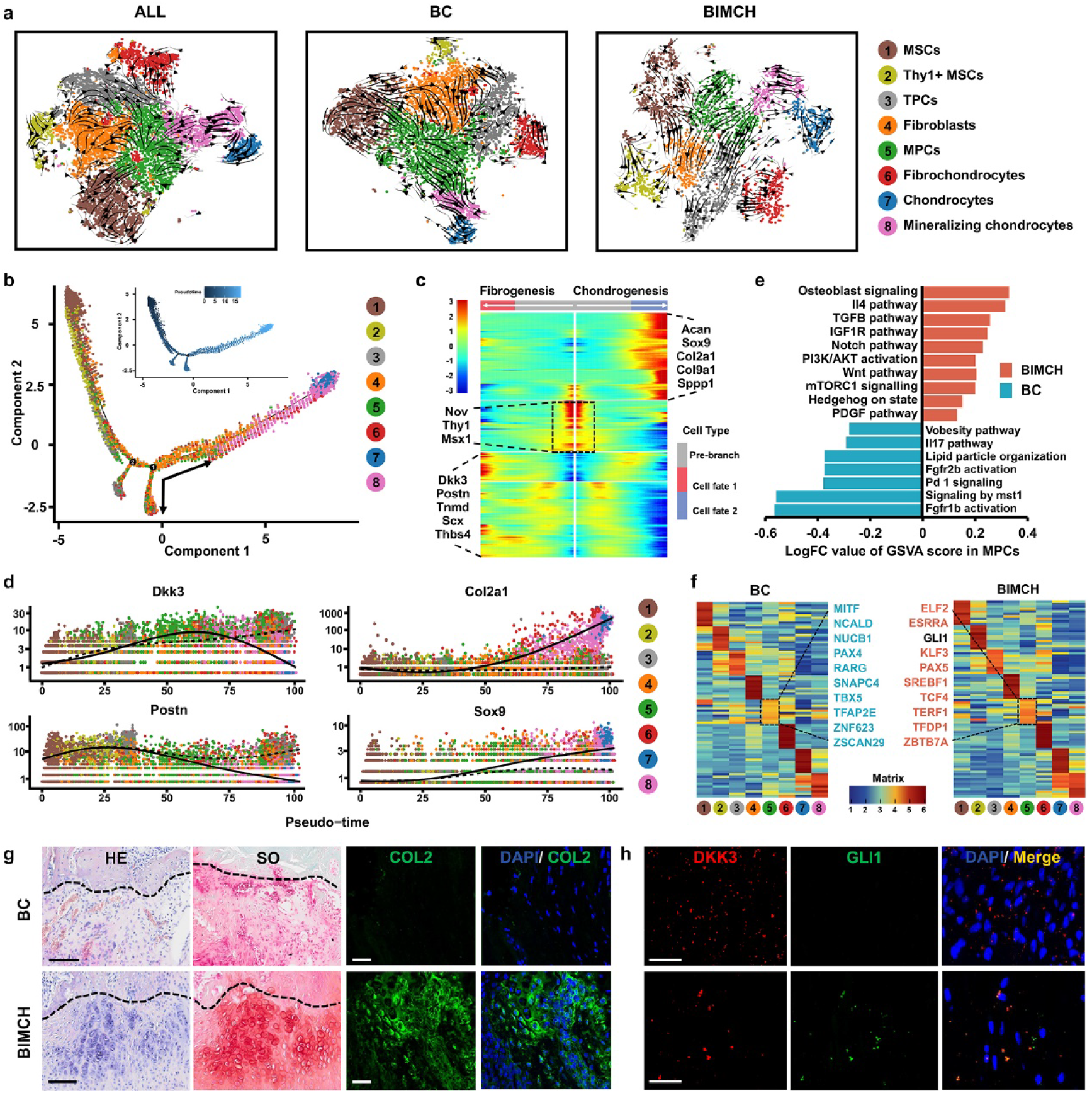
In situ expression of Gli1^+^Dkk3^+^ MPCs promoted efficient TBI regeneration. **a,** RNA velocity-based differentiation trajectory between the BC and BIMCH groups after 2 weeks of defect surgery. **b**, Trajectory of differentiation from MSCs to both fibroblast and chondrocyte lineages predicted by Monocle 2. **c**, Heatmap displaying gene expressions in subsets ordered by pseudotime of the two differentiation trajectories in (b). **d**, Genes (row) clustered, and cells (column) ordered according to pseudotime development. **e**, Pathway enrichment analysis (GSVA) in MPCs from BC and BIMCH groups. LogFC values for each pathway are shown (two-sided unpaired limma-moderated t-test). **f**, Heatmap presenting the AUC score of regulons enriched in entheso-lineage subsets from the two groups. Z-score (row scaling) was calculated. Representative regulons of MPCs are shown between the two heatmaps. **g**, Representative images of H&E (left panel), SO (middle panel), and COL2 (right panel) stained sections of rats treated with the BC and the BIMCH. **h**, Representative images of DKK3 and GLI1 stained sections of rats from the two groups.

Subsequently, we delved into the different mechanisms of MPCs differentiation between the BC and BIMCH groups. Gene Set Variation Analysis (GSVA) revealed that MPCs in the BIMCH group exhibited enrichment in genes associated with the TGF-β pathway, Notch pathway, and Hedgehog signaling, while those in the BC group showed enrichment in genes related to the obesity pathway and lipid particle organization (Fig. 8e). SCENIC analysis demonstrated that MPCs in the BIMCH group were significantly enriched with regulons such as Gli1 (Fig. 8f). Lastly, we substantiated that BIMCH possesses the capability to promote fibrocartilage regeneration as evidenced by SO, SOX9, and COL2 staining (Fig. 8g, Supplementary Fig12a, b). At 2 weeks post-surgery, there was a significantly higher presence of Gli1^+^Dkk3^+^ cells in the BIMCH group compared to the BC group (Fig. 8h, Supplementary Fig12c). By 4 weeks post-surgery, the BIMCH group exhibited a greater abundance of Sox9^+^ and Gli1^+^ cells in the newly formed TBI tissue compared to the BC and NMCH groups (Supplementary Fig13). However, at the 8-week mark post-surgery, the number of Sox9^+^ cells had decreased in the newly formed TBI tissue of the BIMCH group (Supplementary Fig13). In comparison to the BC group, both the NMCH and BIMCH groups displayed higher expression of Sox9^+^ cells in the newly formed tendon tissue. Notably, there was no significant expression of Gli1^+^ cells in the newly formed tendon tissue in any of the three groups at the 8-week time point.

Taken together, we identified that the BIMCH with an ability to direct endogenous cells, Gli1+Dkk3+progenitor cells, to facilitate fibrocartilage regeneration.

## Discussion

In this study, by utilizing FIB-SEM, we discover that within the mineralized region of fibrocartilage, mineral particles do not exclusively align intrafibrillarly or extrafibrillarly; instead, they form a continuous cross-fibrillar phase. Subsequently, through the development of a “floating mineralization” system, we created a bio-inspired mineralization hydrogel consisting of three layers that replicate the hierarchical nano- to micro-scale architecture of TBI. The middle layer, comprising approximately 34 wt% of inorganic content, is particularly noteworthy for its resemblance to the nanostructure of fibrocartilage and its exceptional capacity to induce mineralized fibrochondrogenesis *in vitro*. Through motor function analysis, imaging diagnosis, histological staining, immunofluorescence staining, and biomechanical assessments, we provide evidence that the in situ transplantation of the novel hydrogel achieves synchronized tendon-fibrocartilage-bone regeneration, resulting in a remarkable 68% maximum mechanical recovery at 8 weeks post-operation. Single-cell RNA sequencing analysis reveals that a unique atlas of in situ stem/progenitor cells is generated during the TBI healing *in vivo*. Notably, by single-cell RNA analysis, we show that the BIMCH with an ability to direct endogenous cells, Gli1^+^Dkk3^+^progenitor cells, to facilitate TBI regeneration. Therefore, we have successfully decoded and reconstructed the nanostructure of fibrocartilage, which showed great potential in TBI regeneration.

The design of previous tissue engineering scaffolds has relied on two-dimensional, micrometer resolution structural data, lacking in biomimetic characteristics in high precise manner, thus posing challenges in achieving functional restoration of TBI tissue. Rossetti and colleagues^[25]^ delineate a picture of the fibrous nature of the enthesis at a 3D level using high-resolution microcomputed tomography (µCT). However, in the presence of crystalized CaP, it struggled to analyze the organization and relationship between the mineral and collagen. Recently, the emergence of FIB-SEM technology has provided a novel means to reveal the ultrafine morphological structures of hard tissues. In this study, by utilizing this technique, we have confirmed the spatial relationship between organic and inorganic constituents within fibrocartilage tissue. It provides novel insights into the potential design for high-performance interface materials.

Cells perceive and respond to the biophysical characteristics of biomaterials as they attach to and exert forces on their surrounding synthetic or natural extracellular matrix (ECM)^[3]^. Zhu et al.^[19]^ discovered that there was an opposing pattern of tenogenesis and osteogenesis from the unmineralized region to the highly mineralized region. The expression of *Scx* was predominant in the top unmineralized region, while the expression of *Runx2* was higher than *Scx* in the bottom mineralized region. Similarly, in our observations, when stem cells were seeded onto three distinct materials, a discernible trend emerged where an increase in inorganic constituent content correlated with a reduction in tendon phenotypes and an increasing expression of bone phenotypes. Interestingly, it is noteworthy that in comparison to the fully mineralized collagen, the stem cells cultured on the partially mineralized collagen exhibited the higher expression of chondrogenic phenotypes. As widely accepted, scaffolds containing minerals are effective in inducing osteogenesis and bone growth, making them a common choice for bone induction rather than fibrocartilage. Our data suggests that without the use of partially mineralized collagen for fibrocartilage induction, achieving TBI regeneration remains a distant goal. Therefore, we propose a novel theory and technique for TBI injury regeneration.

The interaction between implanted biomaterials and the local stem/progenitor cells in the microenvironment, subsequently guiding tissue towards fibrosis or regeneration. Dickkopf-3 (*Dkk3*) deficiency significantly ameliorated tubular damage and kidney fibrosis^[26, 27]^. In another research, Arnold et al. discovered that *Dkk3* deficiency promotes the multipotent differentiation potential of induced pluripotent stem cells (iPSCs) *in vitro*, while the absence of *Dkk3 in vivo* reduces liver and pancreatic injury by inhibiting the Wnt signaling pathway and activating the Hedgehog signaling pathway^[28]^. In this study, we observed that, compared to the BC group, the proportion of Dkk3^+^MPCs in the BIMCH group significantly decreased. This may explain the following phenomena: in the BC group, high expression of *Dkk3* promotes tissue fibrosis, while in the BIMCH group, low expression of *Dkk3* initiates the differentiation of MPCs towards other cell lineage, and promotes tissue regeneration. Furthermore, recent research has unveiled a direct involvement of Gli1^+^ progenitor cells in the development of BTI and the repair of associated injuries^[29]^. The formation of the TBI emanates from a distinct reservoir of Gli1^+^ progenitor cells activated by the hedgehog signaling pathway, which then undergo differentiation to generate mineralized fibrocartilage^[30, 31]^. Fang et al found that transplantation of Gli1-lineage cells to mouse enthesis injuries improved healing^[32]^. We also found that MPCs subpopulations from the BIMCH group, relative to the BC group, had their Hedgehog signaling pathway activated, as evidenced by the expression of GLI1 protein. At 2 and 4 weeks post-surgery, immunofluorescence staining revealed a significant increase in Gli+ cells near the newly formed tissue in the BIMCH group compared to the BC group. Therefore, BIMCH can locally activate Dkk3^+^Gli1^+^ progenitor cells, promote their differentiation into chondrocytes, and consequently facilitate the repair of TBI injuries.

## Acknowledgements

The authors are grateful to the Core Facilities of Zhejiang University School of Medicine, Center of Cryo-electron Microscopy of Zhejiang University, and Analysis Center of Agrobiology and Environment Sciences of Zhejiang University for their technical assistance. This work was supported by the NSFC grants (3227100011, 81972099, 82072463, 81871764) and Zhejiang Provincial Natural Science Foundation of China (LZ22H060002).

