## Supplementary figures for "Bio-inspired mineralization collagen induce fibrocartilage regeneration after tendon-bone injury by activating Gli1+Dkk3+ progenitor cells"

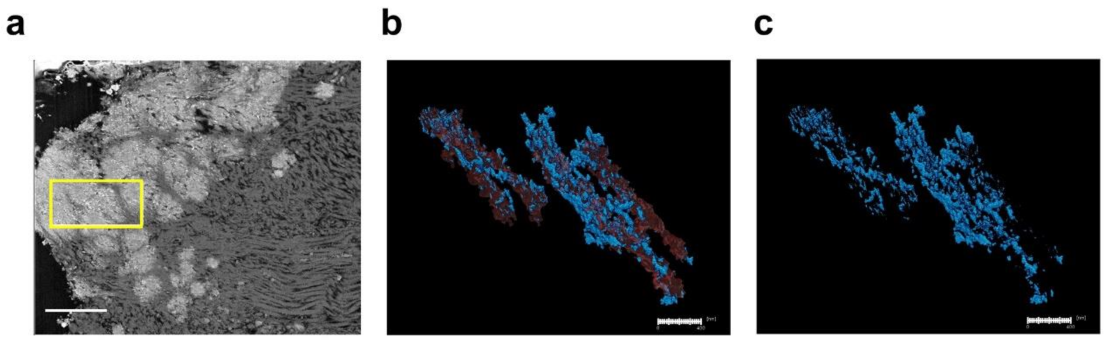


**Figure S1. The high-resolution and three-dimensional images of the bone. a**, the selected area (yellow box) with a scale bar of 2 μm**. b**, The morphology and organization of crystals within the bone are displayed as translucent fibrils (i, scale bar: 400 nm). **c**, only the mineral deposits are displayed with a scale bar of 300 nm.

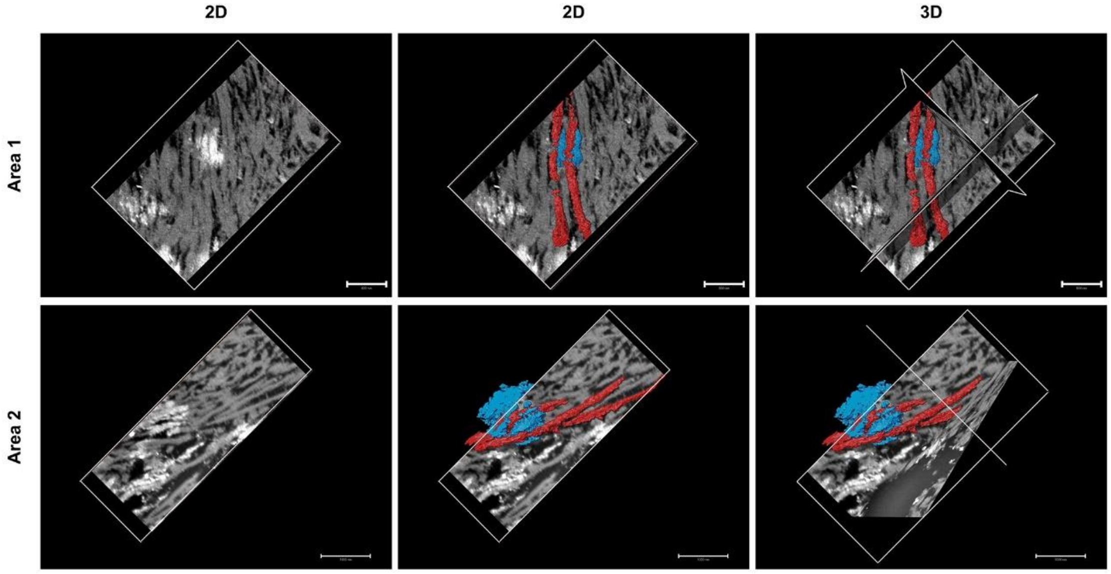


**Figure S2. The two-dimensional (2D) and three-dimensional (3D) localization maps of the selected area 1 and 2.**

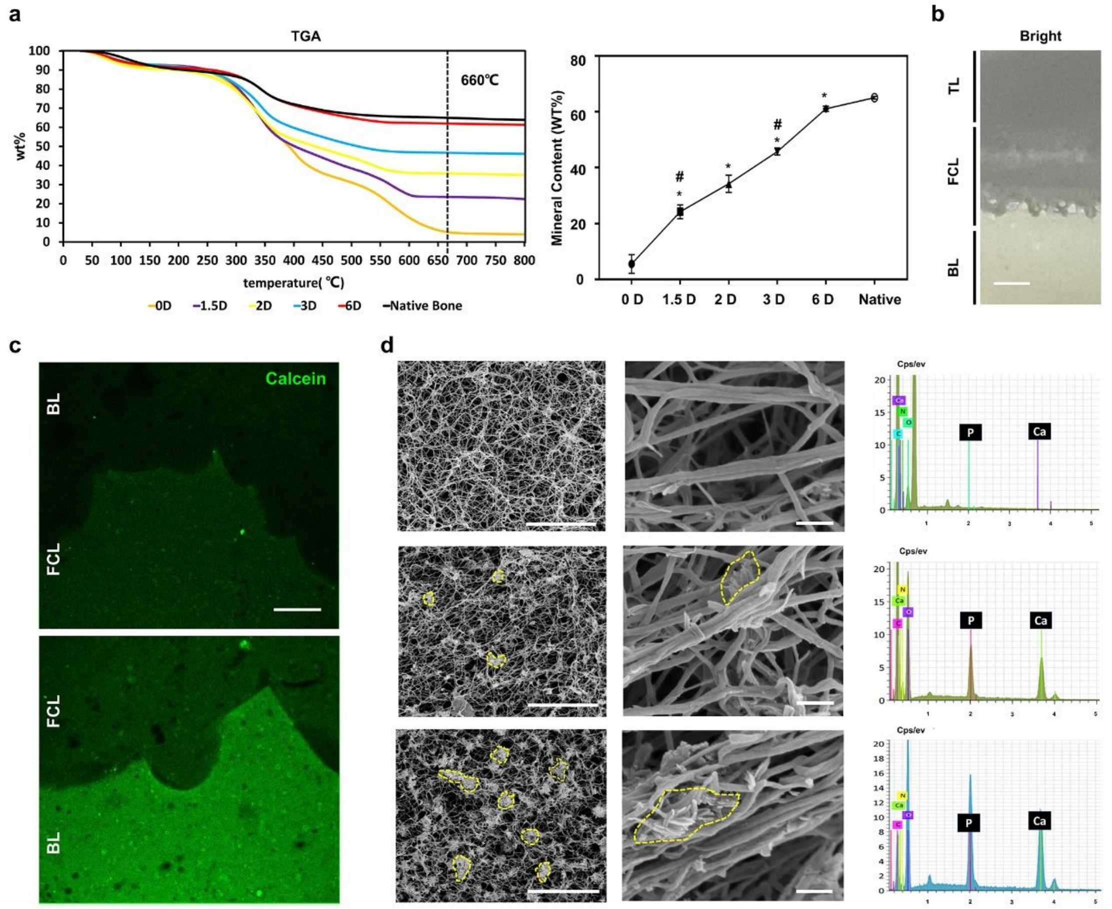


**Figure S3. The bio-inspired tendon–bone interface with gradient nanostructure. a,** Thermogravimetric analysis (TGA) curve (left) and inorganic content (right) of the gel mineralized for different days (D). (n= 3 samples per group). *Comparison with the gel mineralized for 0 days, $P < 0.05$. #Comparison with the gel mineralized for 2 days, $P < 0.05$. **b**, Optical images of the gel demonstrate continuous, interlocked and three-layer structure. Scale bar: 1.5 mm. **c**, Calcein staining images of the gel demonstrate continuous, interlocked and three-layer structure. Scale bar: 200 μm. **d**, SEM showed the porous structure (left panel, Scale bar: 5 μm) and surface roughness (middle panel, scale bar: 400 nm) of the three layers. EDX spectra corresponding to the left panel confirm the relative proportion of Ca and P. The inorganic crystallization was indicated by the yellow dashed line.

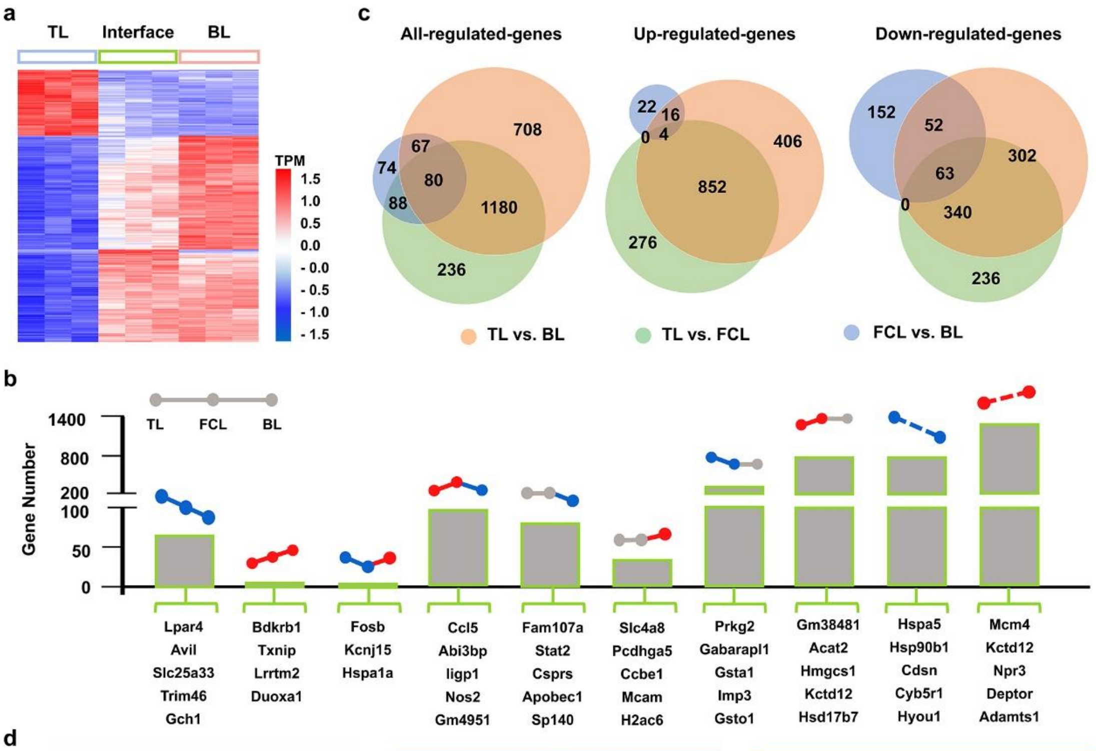


| | TL vs. FCL | NES | FCL vs. BL | NES | TL vs. BL | NES |
|---|---|---|---|---|---|---|
| | GO term | | GO term | | GO term | |
| Up | cholesterol biosynthetic process | 2.24 | DNA replication initiation | 2.50 | cholesterol biosynthetic process | 2.13 |
| | sterol biosynthetic process | 2.00 | mitotic sister chromatid segregation | 2.48 | DNA replication initiation | 2.06 |
| | peptidyl-tyrosine phosphorylation | 1.96 | DNA replication | 2.35 | mitotic sister chromatid segregation | 2.04 |
| | defense response | 1.94 | cell division | 2.34 | signal transduction | 2.03 |
| | steroid biosynthetic process | 1.92 | mitotic cytokinesis | 2.26 | DNA replication | 2.02 |
| Down | ribosomal large subunit biogenesis | -2.95 | cellular response to interferon-beta | -2.98 | response to unfolded protein | -2.75 |
| | cytoplasmic translation | -2.69 | defensive response to virus | -2.73 | ribosomal large subunit biogenesis | -2.71 |
| | ribosomal large subunit assembly | -2.42 | T cell mediated cytotoxicity | -2.60 | cellular amino acid biosynthetic process | -2.67 |
| | ribosome biogenesis | -2.26 | response to bacterium | -2.59 | cytoplasmic translation | -2.66 |
| | response to unfolded protein | -2.26 | antigen processing and presentation of peptide antigen via MHC class I | -2.53 | endoplasmic reticulum unfolded protein response | -2.63 |

**Figure S4. BIMCH induced MSCs toward TBI cell lineage differentiation *in vitro*. a**, Heatmaps displaying transcript per million (TPM) values of the top 500 differentially expressed genes (DEGs, P < 0.05) in MSCs after a 3-day exposure to the TL layer, FCL layer, and BL layers, respectively. Red indicates upregulated genes, while blue represents downregulated genes. **b**, Classification of DEGs (P < 0.05, > 2-fold changes). The bar chart illustrates the number of DEGs corresponding to each stiffness response profile. Gray color-coding denotes a stiffness range with no statistically significant differential expression, while green and red color-coding indicate positive and negative, statistically significant differential expression, respectively. **c**, Venn Diagram depicting all differentially expressed genes (p<0.05, 2-fold change compared to controls). **d**, Top 5 Gene Ontology (GO) terms related to biological processes.

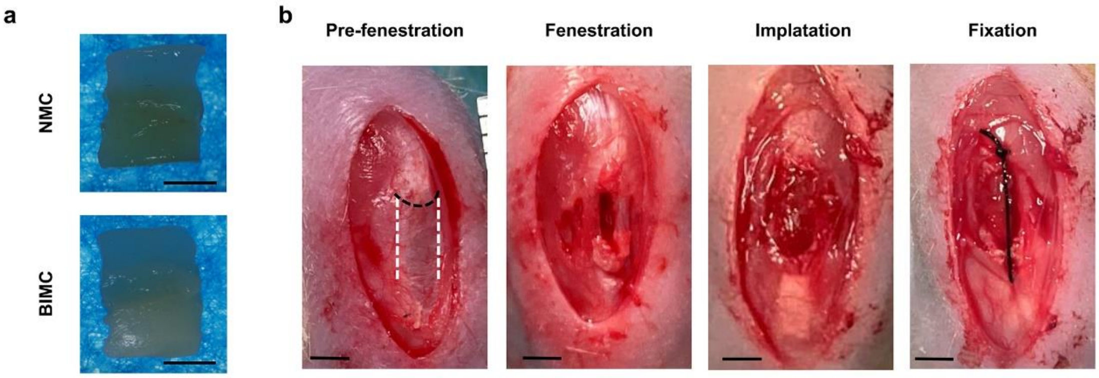


**Figure S5. *In vivo* implantation of the material.** **a**, The appearance of the implanted material *in vivo*, scale: 1 mm. **b**, The patellar-to-tendon interface fenestration model.

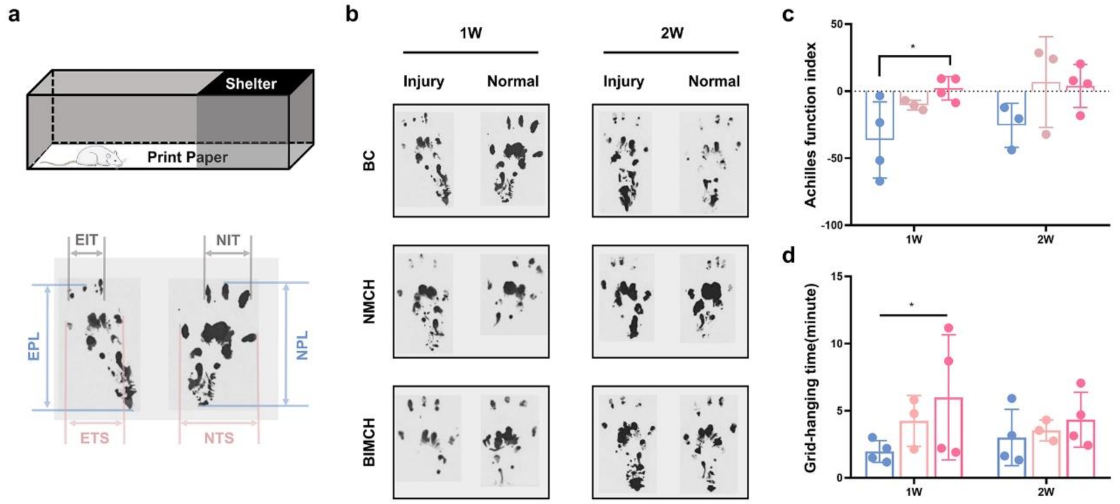


**Figure S6. Functional evaluation of tendon-bone tissue regeneration. a**, Schematic diagram of the footprint experiment. Representative images (**b**) and quantitative analysis results (c) of the footprint experiment at 1 and 2 weeks post-surgery (n = 3-4). print length (PL), toe spreading (TS), intermediary toes (IT), normal (N), experiment (E). **d**, Quantitative analysis results of the grid-hanging experiment (n = 3-4). W: Week.

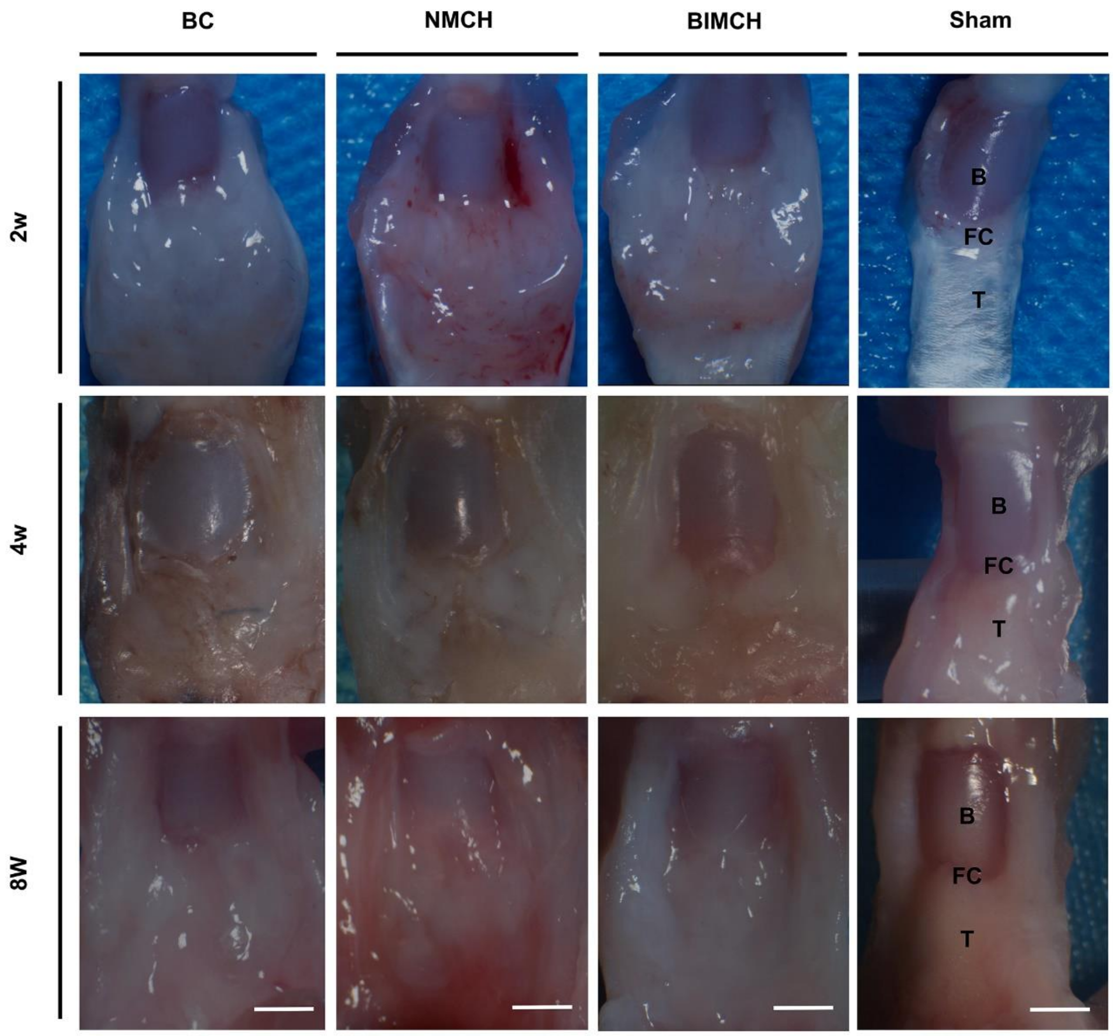


**Figure S7. Macroscopic evaluation of TBI Regeneration (posterior aspect of the patella).** B: bone, FC: fibrocartilage, T: tendon, W: week. n = 5, scale bar: 2 mm.

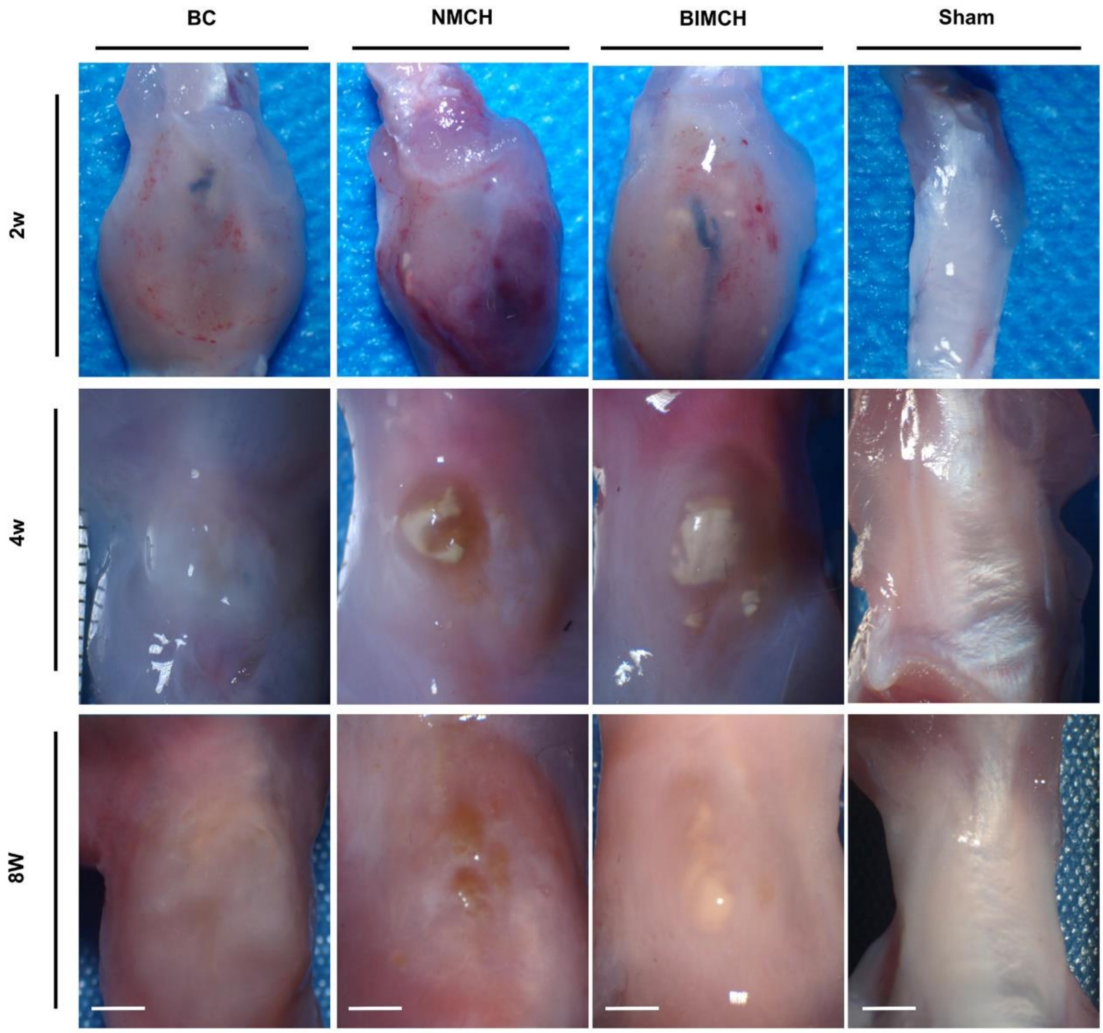


**Figure S8. Macroscopic evaluation of TBI Regeneration (anterior aspect of the patella).** n = 5, scale bar: 2 mm.

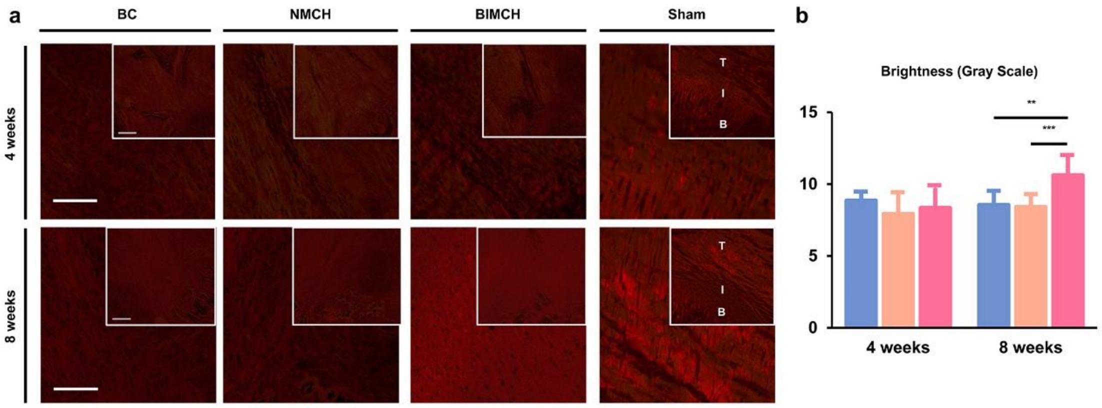


**Figure S9. Representative images of picrosirius red–stained sections of rats treated with the BC, NMCH, and the BIMCH (n = 5).** T, tendon; I, interface; B, bone. Scale bars: 100 µm, 200 µm (inset).

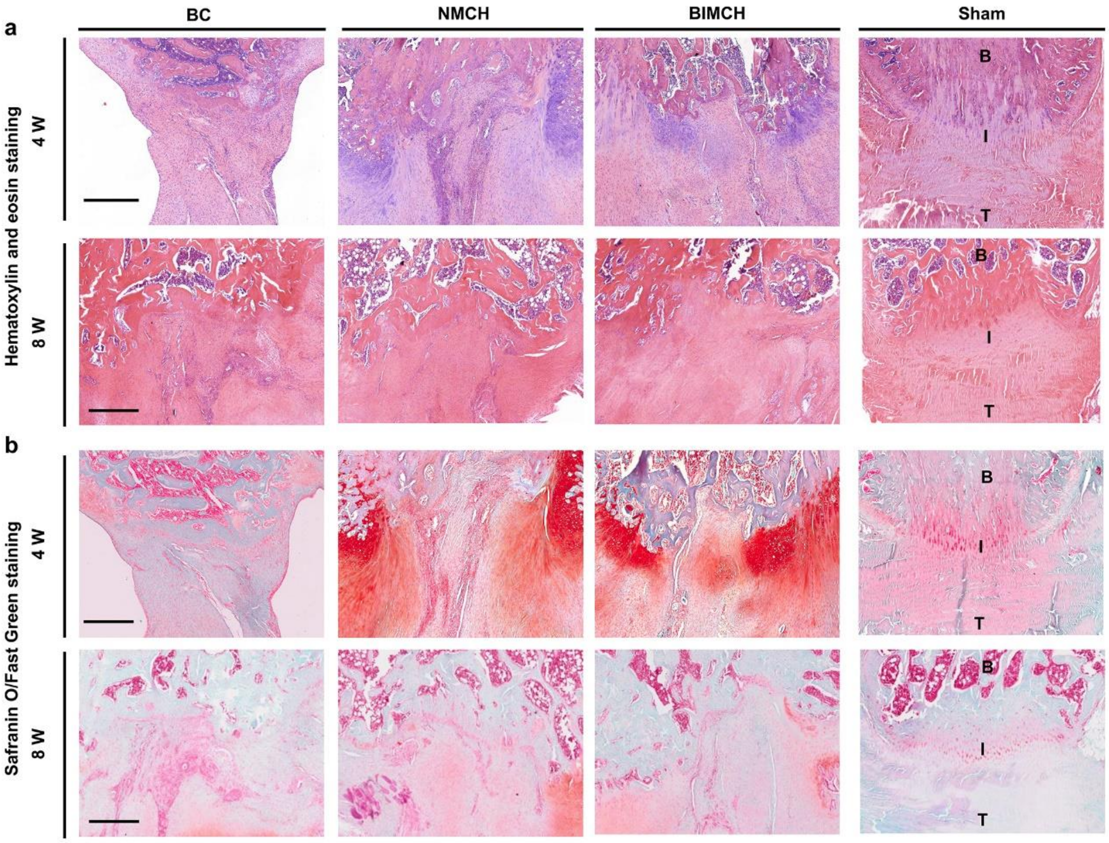


**Figure S10. Morphological analysis of the newly formed tendon-bone interface tissue after treatment.** Representative images of H&E and SO stained sections from rats treated with BC, NMCH, and the BIMCH (n = 5). T, tendon; I, interface; B, bone. Scale bars: 500 μm.

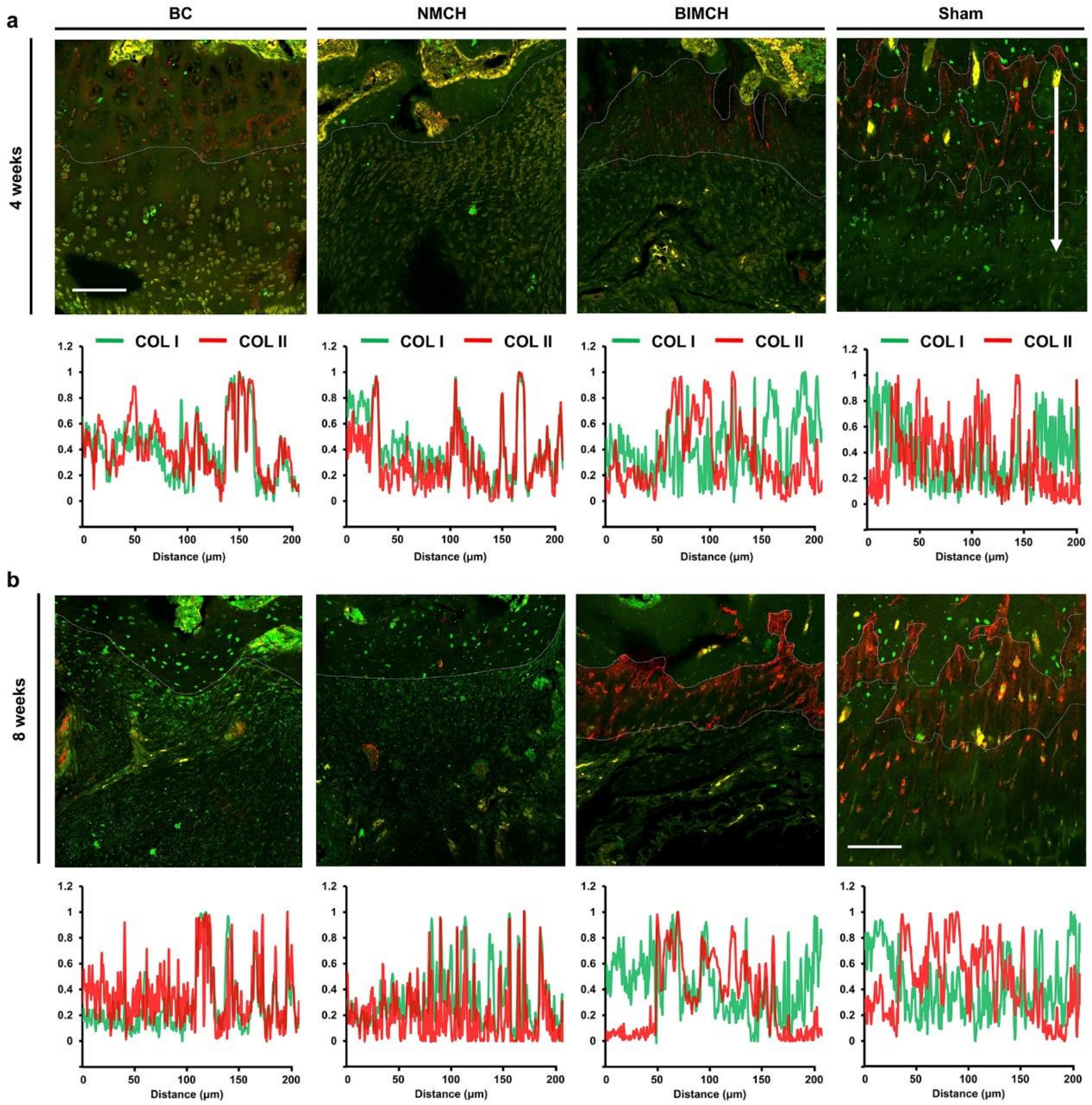


**Figure S11. Collagen composition analysis of the newly formed tendon-bone interface tissue at 4 weeks (a) and 8 weeks (b) post-operation (n=3).** The white dotted lines delineate the boundaries of different tissue types. Fluorescence intensities are plotted over distance across the insertion (indicated by the long arrow in the confocal image of the sham group at 4 weeks post-operation, averaged over a width of 200 μm). Scale bar: 100 μm.

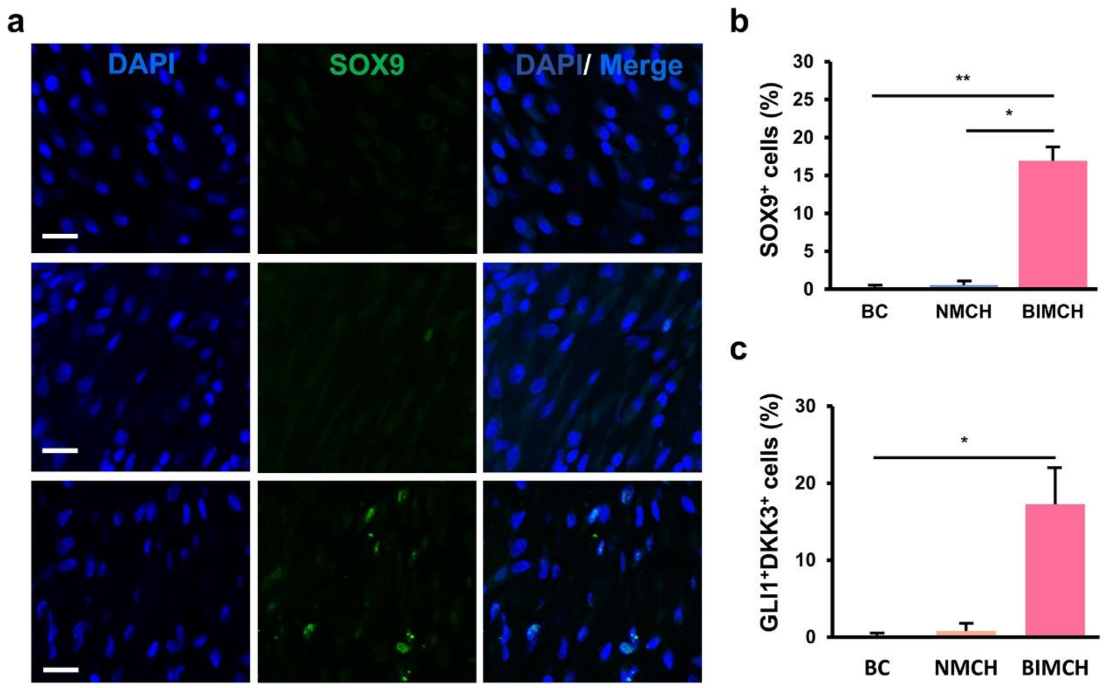


**Figure S12. At 2 weeks post-surgery, the compositional differences in the newly formed TBI tissues among the three groups were evaluated (n = 4). a-b,** The differential expression of $Sox9^{+}$ cells in the newly formed TBI tissues of the three groups (a) and the quantitative analysis results (b) are shown (scale bar: 30 μm). **c**, The differential expression of $Gli1^{+}Dkk3^{+}$ cells in the newly formed TBI tissues of the three groups.

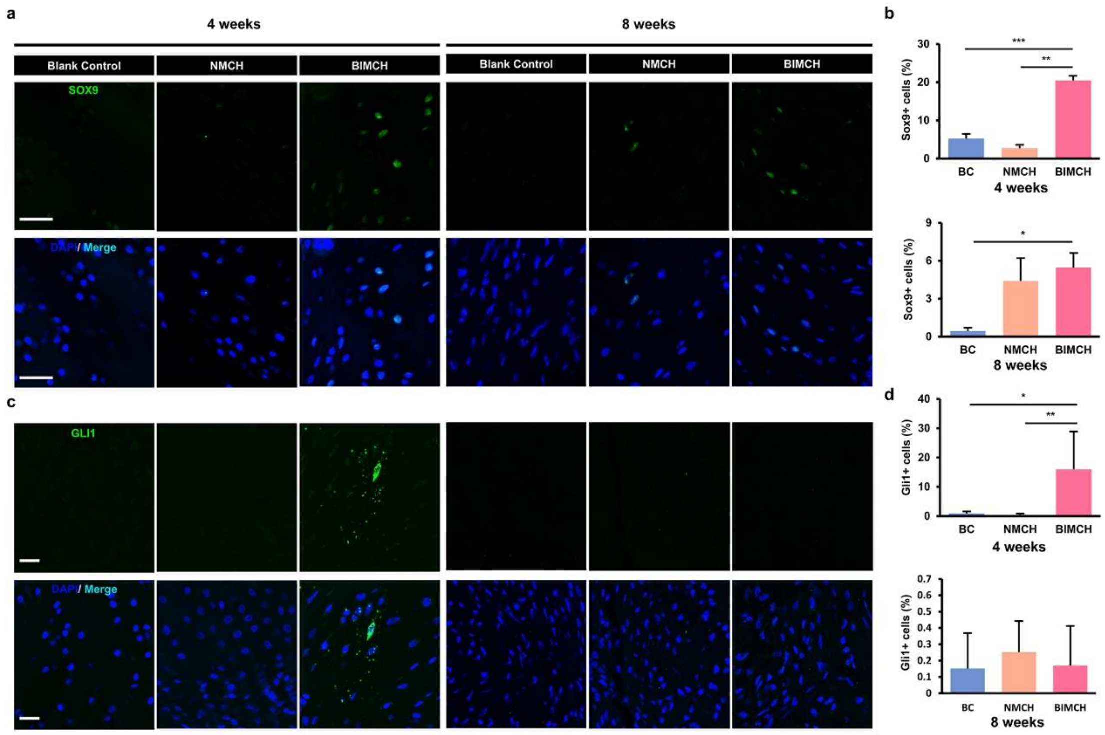


**Figure S13. At 4 and 8 weeks post-surgery, the compositional differences in the newly formed TBI tissues among the three groups were evaluated (n = 4). a-b,** The differential expression of $Sox9^+$ cells in the newly formed TBI tissues of the three groups (a) and the quantitative analysis results (b) are shown (scale bar: 30 μm). **c-d**, The differential expression of $Gli1^+$ cells in the newly formed TBI tissues of the three groups (c) and the quantitative analysis results (d) are shown (scale bar: 20 μm).

### Materials and methods

**Sample preparation for structure analysis.** Achilles tendon–bone samples were extracted from porcine legs obtained from a local abattoir. The pigs were 6 months old at the time of slaughter. The samples were then sliced sagittally into 2 mm sections using an oscillating saw. These sample slices were subsequently fixed in 2.5% glutaraldehyde at 4°C overnight. For transmission electron microscopy (TEM) and scanning electron microscopy (SEM), the sample slices were cryo-sectioned into 100-μm-thick sections.

**Focused Ion Beam-Scanning Electron Microscopy.** After the initial fixation, the samples were rinsed with 0.1 M PBS three times and then fixed in osmic acid at 4°C for 1 hour. Subsequently, the samples underwent dehydration through a graded ethanol series and were embedded in acrylic resin. Sections were stained with uranyl acetate and lead citrate. A scanning electron microscope (Thermo Fisher, Teneo VS) equipped with an ultramicrotome in its specimen chamber for trimming the resin blocks and acquiring electron microscopic images of the samples simultaneously was utilized. Imaging was conducted using a dual-beam scanning electron microscopy system (Thermo Fisher, FIB Helios G3 UC). The data collection procedure operated in serial-surface view mode with a slice thickness of 5 nm at 30 keV and 0.79 nA. Each serial face was imaged at 2 kV acceleration voltage and a current of 0.2 nA in backscatter mode (BSE) with an ICD detector. The image store resolution was set to 3072×2048 pixels with a dwell time of 15 μs per pixel and 4.25 nm per pixel. A total of 695 slices were acquired, capturing one entire sensilla in the image stack with a depth of 3.48 μm. The resulting image stacks were segmented based on their structural features and reconstructed using Amira 3D (v 6.5; Thermo Fisher and Zuse Institute).

**Collagen Extraction.** Porcine tendons were collected postmortem. All tools and vessels were autoclaved before use. Deionized water was employed, and all solutions were pre-filtered through 0.22 μm Millipore filters. To summarize, porcine tendons were initially minced into small pieces and immersed in a solution containing 0.05 M Tris, 0.02 M EDTA, and 0.5 M NaCl for 24 hours at 4°C. While being stirred, the tendons were dissolved in a solution consisting of 0.5 M acetic acid and pepsin (100 mg per 1g of tendons) for 3 days at 4°C. The resulting viscous solution was then centrifuged at 4000 rpm and 4°C for 25 minutes. The supernatant was collected and reprecipitated by the addition of 0.9 M NaCl, allowing reprecipitation to occur overnight at 4°C with continuous stirring. The collagen was subsequently centrifuged for 25 minutes at 4000 rpm and 4°C to collect the precipitated collagen (pellet). The precipitate was redissolved in 0.5 M acetic acid overnight. The resulting solution was dialyzed over 3 days at 4°C, with regular changes of the dialysate. The collagen solution was then transferred to 50 mL conical tubes and lyophilized. The lyophilized collagen was stored at -20°C and redissolved in acetic acid (pH=3) to a concentration of 10 mg mL-1 for subsequent tests.

**Collagen hydrogel manufacturing and mineralization.** A total of 100 μL of Collagen-I solution was placed either into cylindrical molds (with a diameter of 4 mm and a height of 8 mm) or in 24-well plates. The self-assembly process of collagen molecules into fibrils was initiated by neutralization with ammonia vapor, which was carried out for 4 hours. The hydrogel formed in the cylindrical molds could be extracted and transferred into 6-well plates for subsequent tests. The hydrogel was gently rinsed with deionized water until a neutral pH was achieved. The collagen hydrogel was further cross-linked using a 0.03 wt% glutaraldehyde solution for 2 hours, followed by another gentle rinse with deionized water. Mineralization was then performed using a mineralization solution containing 1.67 mM CaCl2, 240 μg mL-1 P-Asp, 9.5 mM Na2HPO4, and 150 mM NaCl. To prepare the mineralization solution, a specific volume of calcium solution (3.34 mM CaCl2, labeled as A1) was dripped into a Petri dish. Then, a designated volume of 10 mg mL-1 p-Asp stock solution was added to the calcium solution and thoroughly mixed (referred to as solution A2). Subsequently, a phosphate solution (with an equal volume to the calcium solution, consisting of 19 mM Na2HPO4 and 300 mM NaCl) was added to solution A2 and mixed thoroughly. The gels were subjected to mineralization at 37 °C under 100% relative humidity for a specified duration. After the mineralization process, the gels were extensively washed to remove any residual salts.

**Gradient bio-inspired mineralization collagen hydrogel manufacturing.** A 100 μL volume of Collagen-I solution was placed into cylindrical molds. After self-assembly and cross-linking, the first layer of BIMCH was subjected to mineralization for 4 days. Subsequently, 50 μL of acid-soluble collagen was gelled in contact with the first layer. After self-assembly and cross-linking of this additional layer, the two layers of BIMCH were mineralized for 2 days. Then, 100 μL of acid-soluble collagen was gelled in contact with the second layer. After self-assembly and cross-linking of this final layer, BIMCH was obtained. The production method for NMCH is identical to that of BIMCH, with the exception that NMCH is not subjected to the mineralization process.

**Scanning electron microscopy of the hydrogel.** For SEM analysis, samples were fixed with 2.5% glutaraldehyde for 1 h at room temperature, washed in distilled water, and subjected to a series of ethanol dehydration steps for 10 min each. Subsequently, the samples were critical point dried, sputter coated with gold/palladium, and observed under SEM (Hitachi SU-8010). EDXS spectra were collected using SU8010 equipped with an energy dispersive x-ray spectrometer (Model 550i, IXRF Systems).

**Transmission electron microscopy of the hydrogel.** TEM was performed with a Hitachi HT-7700 (Japan) operating at 120 kV. Both mineralized and non-mineralized hydrogels were minced with a double-edge razor blade and were immersed in ice-cold 0.1 M ammonium bicarbonate (pH 7.8). The minced hydrogels were then exposed to the tissue homogenizer operated at 65Hz until no visible fragments remained. The homogenate (3μL) was then dripped onto 300-mesh carbon-and-formvar-coated Nickel TEM grids. Non-mineralized hydrogels on TEM grids were stained with uranyl acetate for 15 s.

**Thermogravimetric analysis.** TGA was performed using a TA Instrument (Mettler Toledo Corp., Switzerland). Samples were dried at 37°C for 3 days. The sample temperature was then heated under air conditions from room temperature to 800°C. The mass at 660°C was considered as the total inorganic mass.

**FTIR analysis.** FTIR (IRAffinity-1, Shimadzu, Japan) was performed with 30 scans at 4 $cm^{-1}$ resolution from 4000 to 400 $cm^{-1}$. T The background was established using blank KBr plates.

**Young's modulus of hydrogels.** Young's modulus of the hydrogels was measured using a nanoindenter (Piuma Chiaro, Optics11, The Netherlands). The hydrogels were affixed to the bottom of a Petri dish and submerged in deionized water. A spherical nano-indentation probe with a tip radius of 46.5 μm and a stiffness of 0.5 N m^-1 was employed. Load-indentation data were recorded as the probe made contact with the material's surface. The indentation depth was set to 10 μm, and the loading and unloading periods were each set to 2 seconds. The Effective Young's modulus was determined based on the load-indentation curve using Optics11.

**Stochastic Optical Reconstruction Microscopy (STORM) was utilized to acquire 3D images of the mineralized collagen fibrils' structure.** The stochastic optical reconstruction microscopy (STORM) technique[1, 2] enables subdiffraction-level accuracy for confirming the intra- and extrafibrillar mineralization of collagen-I fibrils. Following a 2-day mineralization period, the collagen hydrogel was incubated overnight at 4°C with a rabbit anti-mouse antibody targeting collagen-I (abcam, ab34710). Subsequently, the hydrogel underwent a 1.5-hour incubation with fluorescein-conjugated secondary antibodies. Finally, the mineralized hydrogel was labeled with calcein at a concentration of 10 × 10^-6 M for 40 minutes. All STORM imaging experiments were conducted using a Nikon Ti-E inverted optical microscope, and the resulting movies and images were analyzed with Nikon NIS-Elements AR software.

**Cell culture**. C3H10T1/2 cells (mouse multipotent MSC line) were obtained from the Cell Bank of the Chinese Academy of Sciences (Shanghai, China, http://www.cellbank.org.cn/). Cells were cultured in Dulbecco's modified Eagle's medium (DMEM, low glucose; Gibco) with 1% penicillin-streptomycin and 10% fetal bovine serum.

**Cell viability.** Cell viability was assessed by the live/dead cell staining (Yeasen Biotechnology) at 1, 4, and 7 days, according to the manufacturer's instructions. Briefly, the cells were washed 2-3 times with Assay Buffer. Next, the cells were incubated with solution containing 2 μM Calcein-AM and 4.5 μM PI at 37 °C for 30 min. Images were captured using confocal microscope (Olympus).

**Cell morphology.** Cell phenotype assessment was conducted after 7 days of culture on the TL layer, FCL layer, and BL layer. The cells were detached and plated on a well plate. Subsequently, they were fixed in 4% paraformaldehyde for 15 minutes, permeabilized with 1% Triton X-100 in PBS for 10 minutes, and blocked with 1% BSA for 30 minutes. The cells were then exposed to primary antibodies overnight at 4 °C. The primary antibodies employed were anti-SOX9 (Abcam) and anti-RUNX2 (Abcam). Following a PBS rinse, the cells were treated with goat anti-rabbit secondary antibodies labeled with Alexa Fluor 488 (Invitrogen). To visualize cell nuclei, DAPI was used. The images were captured using a confocal microscope (Olympus).

**Cell phenotype.** To assess cell phenotype, cells were cultured on the TL layer, FCL layer, and BL layer for 7 days. These cells were detached and plated onto a well plate. Afterward, the cells were fixed with 4% paraformaldehyde for 15 minutes, permeabilized using 1% Triton X-100 in PBS for 10 minutes, and then blocked with 1% BSA for 30 minutes. Subsequently, the cells were subjected to overnight incubation at 4 °C with primary antibodies, namely anti-SOX9 (Abcam) and anti-RUNX2 (Abcam). Following PBS washing, the cells were exposed to a secondary antibody, goat anti-rabbit conjugated with Alexa Fluor 488 (Invitrogen). To visualize cell nuclei, DAPI was utilized. The resulting images were captured using a confocal microscope (Olympus).

**Bulk RNA-seq.** For bulk RNA-seq analysis, cells were cultured on the TL layer, FCL layer, and BL layer for 3 days. RNA was extracted using the RNA-Quit Purification Kit from ESScience Biotech. After confirming RNA quality, reverse transcription was carried out to generate a cDNA library (performed by BGI-Shenzhen, China) for subsequent sequencing. To conduct functional analysis, differential gene ontology (GO) or pathway analyses were performed on the three groups using the Gene Set Enrichment Analysis (GSEA) tool from the GSEA package (version 4.1.0). The analysis utilized curated gene sets, specifically the hallmark gene sets from the MSigDB databases (https://www.gsea-msigdb.org/gsea/msigdb). To attribute functional differences and activity estimates to the groups, we employed the non-parametric and unsupervised GSVA (Gene Set Variation Analysis) algorithm, available in the GSVA package (version 1.38.2).

**Surgical procedure.** Eight-week-old male SD rats, obtained from SLAC Laboratory Animal in Shanghai, China, were randomly divided into three groups: the BIMCH group, NMCH group, and BC group. All procedures were conducted following approved protocols (ZJU20210029) in accordance with the guidelines set by the animal experimental center of Zhejiang University, China. Under general anesthesia, a skin incision was made between the upper edge of the patella and the tibial tubercle. Subcutaneous tissue was carefully separated to expose the patella and tendon. A fenestration model, measuring 1.5 mm in width and 5 mm in length, was created at the patellar-tendon interface tissue, including the distal 3 mm of the patella along with its connected fibrocartilage layer and 2 mm of the patellar tendon. Following this, BIMCH or NMCH grafts were cut into appropriate dimensions (1.5 mm in width, 1 mm in thickness, and 5 mm in length) and placed within the fenestration site. In the case of the defect group, no graft implantation was performed. Rats were euthanized at 2, 4, and 8 weeks postoperatively, and specimens of the quadriceps-patella-patellar tendon-tibia complex (QPPTC) were collected and observed for further analysis.

**Footprint experiment.** We employed a method described in a prior study[3]. Achilles Functional Index (AFI) values for healthy rats typically approach zero, with a more negative value indicating greater impairment in motility. Briefly, a restrictive roadway (80 cm long, 9 cm wide, 20 cm height) was covered with a white paper. After their hind paws were evenly dipped with black ink, the rats were allowed to walk freely, printing black footprints on the white paper. To quantify the AFI of the rats, footprints were scanned. We defined and acquired related footprints' parameters including print length (PL), toe spreading length (defined as the distance between the first and fifth toes, TS), and intermediary toe spreading length (defined as the distance between the second and fourth toes, IT). Then, according to the difference between the normal (N) and the experimental values (E), three footprints dimension factors including print length factor (PLF), toe spreading length factor (TSF), intermediary toe spreading length factor (ITF) could be acquired by use of the following equations:

$$PLF = NPL\text{-}EPL/EPL \quad (1)$$

$$TSF = (ETS\text{-}NTS)/NTS \quad (2)$$

$$ITF = EIT - NIT/NIT \quad (3)$$

Finally, the AFI was calculated according to an established equation[3]

$$AFI = 74 \times (PLF) + 161 \times (TSF) + 48 \times (ITF) - 5$$

**Inverted grid test.** Rats were placed in the center of the wire grid system, and the grid was rotated to an inverted position over 2 s, with the mouse head declining first. The grid was held steadily 40–50 cm above a padded surface. The grid-hanging time capacity was determined by the time the time the rats spent hanging on the grid.

**Micro-CT imaging and analysis.** The QPPTC specimens from the rats were fixed in 4% paraformaldehyde and subjected to analysis using micro-CT (Skyscan 1172). The micro-CT scanner operated at a voltage of 80 kV with a resolution of 18 μm per pixel.

Subsequently, the obtained images were reconstructed and analyzed to determine the patellar volume. The reconstruction and analysis processes were conducted using software tools such as NRecon, CTAn, and CTVol.

**Histology and immunofluorescence staining.** The tissue specimens were initially fixed in a 4% paraformaldehyde solution, followed by a thorough washing with running water. Subsequently, they were dehydrated through a graded series of ethanol, vitrified using dimethylbenzene, and ultimately embedded in paraffin. Sections with a thickness of 7 μm were obtained from the paraffin-embedded specimens. To prepare these sections for analysis, they were deparaffinized using xylene and then rehydrated through a gradient of ethanol solutions. Staining was performed using standard H&E staining, SO staining, and Sirius Red staining protocols. The organization of collagen fibers was evaluated using polarizing microscopy. Furthermore, histological scores were calculated based on the results obtained from H&E staining. For immunofluorescence (IF) analysis, sections were subjected to overnight incubation at 4°C with primary antibodies. Specifically, the primary antibodies used in this study included anti-COL1 (abcam, ab34710), anti-COL2 (Pretech, 28459-1-AP), anti-SOX9 (abcam, ab185230), and anti-GLI1 (R&D Systems, AF3455). Following the primary antibody incubation, the sections were exposed to fluorescein-conjugated secondary antibodies for 1.5 hours and subsequently observed under a confocal fluorescence microscope (Nikon).

**Mechanical testing.** Mechanical testing was performed using a tension/compression system with the Fast-Track software (Instron). The hind limbs were collected and all soft tissue spanning the knee, except for the center of the patellar and tendon, were transected. Measurement of the tendon cross-sectional area was performed, and the QPPTC was then rigidly fixed to custom-made clamps. After applying a preload of 0.1 N, each FPTC underwent preconditioning by cyclic elongation of 0 and 0.5 mm for 20 cycles at 5 mm/min. This was followed by a load to-failure test at an elongation rate of 5 mm/min. The structural properties of the FPTC were represented by stiffness (N/mm), failure force (N), energy absorbed at failure (mJ), modulus (MPa) and stress at failure (MPa).

**Preparation of single-cell suspensions from rat patella-tendon complex.** At 2 weeks post-surgery, the rats were anesthetized to initiate cardiac perfusion with phosphate-buffered saline (PBS). Following euthanasia, patella-patellar tendon specimens were carefully extracted from the rats and rinsed with PBS. These tissues were finely minced using razor blades, washed repeatedly with PBS, and subsequently subjected to enzymatic digestion (comprising 0.1% type I collagenase and 0.1% dispase) at 37 °C for 40 minutes. The resulting mixture was then passed through a 70 μm nylon mesh to obtain a single-cell suspension. Subsequently, mRNA libraries were prepared using the GEXSCOPE® Single Cell RNA Library Kits and subjected to sequencing.

**Processing of scRNA-seq data.** For quality control, we retained cells that exhibited the following criteria: detection of over 300 genes, a read count of at least 500, and mitochondrial gene expression below 20%. After applying these filters, we obtained data from a total of 15,866 cells, with 9,619 cells in the BC group and 6,247 cells in the BIMCH group. We followed the Seurat pipeline for further analysis, which included log-normalization and scaling of the data. To visualize cell heterogeneity in lower dimensions, we employed Uniform Manifold Approximation and Projection (UMAP) as well as t-distributed Stochastic Neighbor Embedding (t-SNE). For single-cell clustering and annotation, after dimensionality reduction, we conducted clustering using the FindClusters function provided by Seurat. Genes that were specifically expressed in each cluster were identified using the Seurat FindAllMarkers function. To determine the biological cell type of each cluster, we performed SingleR analysis, complemented with traditional markers for some known cell types. In order to map the differentiation trajectory in TBI regeneration, we conducted pseudotime analysis using the R package Monocle (version 2.20.0). Specifically, we computed the trajectory of entheso-lineage cells and compared different cell states using the BEAM function from Monocle. RNA velocity analysis was performed using the scvelo python package (version 0.1.25). For gene functional annotation analysis, we carried out GO enrichment analysis for markers of single-cell clusters using the clusterProfiler package (version 81). The enriched GO terms were filtered using a p-value cutoff of 0.01. To analyze single-cell regulatory networks, we utilized the SCENIC package (version 82) and followed the standard pipeline. The dot plot displays cell-type-specific regulons with the top Regulon Specificity Score (RSS) and their average expression (Z) in the cell subtype. Differential analysis between the GIMCH group and the blank control was conducted using the Wilcoxon rank sum test.

**Statistics analysis.** In all graphs, the data are presented as the mean ± standard deviation. For the experiments involving the comparison of two groups, statistical analysis was performed using two-tailed, unpaired Student's t-test (SPSS 25.0 software). For experiments involving more than two groups, one-way ANOVA with post hoc Tukey's test or Kruskal–Wallis test with post hoc Dunn's test for multiple comparisons was used to identify significant differences. P-value of less than 0.05 is considered significant (significance is denoted as follows: *$P<0.05$, **$P<0.01$ and ***$P<0.001$).
